# A metabolic reconstruction of *Lactobacillus reuteri* JCM 1112 and analysis of its potential as a cell factory

**DOI:** 10.1101/708875

**Authors:** Thordis Kristjansdottir, Elleke F. Bosma, Filipe Branco dos Santos, Emre Özdemir, Markus J. Herrgård, Lucas França, Bruno Sommer Ferreira, Alex T. Nielsen, Steinn Gudmundsson

## Abstract

**Background:** *Lactobacillus reuteri* is a heterofermentative Lactic Acid Bacterium (LAB) that is commonly used for food fermentations and probiotic purposes. Due to its robust properties, it is also increasingly considered for use as a cell factory. It produces several industrially important compounds such as 1,3-propanediol and reuterin natively, but for cell factory purposes, developing improved strategies for engineering and fermentation optimization is crucial. Genome-scale metabolic models can be highly beneficial in guiding rational metabolic engineering. Reconstructing a reliable and a quantitatively accurate metabolic model requires extensive manual curation and incorporation of experimental data.

**Results:** A genome-scale metabolic model of *L. reuteri* JCM 1112^T^ was reconstructed and the resulting model, Lreuteri_530, was validated and tested with experimental data. Several knowledge gaps in the metabolism were identified and resolved during this process, including presence/absence of glycolytic genes. Flux distribution between the two glycolytic pathways, the phosphoketolase and Embden-Meyerhof-Parnas pathways, varies considerably between LAB species and strains. As these pathways result in different energy yields, it is important to include strain-specific utilization of these pathways in the model. We determined experimentally that the Embden-Meyerhof-Parnas pathway carried at most 7% of the total glycolytic flux. Predicted growth rates from Lreuteri_530 were in good agreement with experimentally determined values. To further validate the prediction accuracy of Lreuteri_530, the predicted effects of glycerol addition and *adhE* gene knock-out, which results in impaired ethanol production, were compared to *in vivo* data. Examination of both growth rates and uptake- and secretion rates of the main metabolites in central metabolism demonstrated that the model was able to accurately predict the experimentally observed effects. Lastly, the potential of *L. reuteri* as a cell factory was investigated, resulting in a number of general metabolic engineering strategies.

**Conclusion:** We have constructed a manually curated genome-scale metabolic model of *L. reuteri* JCM 1112^T^ that has been experimentally parameterized and validated and can accurately predict metabolic behavior of this important platform cell factory.

## 1. Introduction

*Lactobacillus reuteri* is a heterofermentative Lactic Acid Bacterium (LAB) that is present in the human gut and is an important probiotic organism (Saulnier et al., 2011). There is an increasing interest in using it as a cell factory for the production of green chemicals and fuels in a biorefinery (Dishisha, Pereyra, Pyo, Britton, & Hatti-Kaul, 2014; Ricci et al., 2015), due to its robustness properties. It has high growth and glycolytic rates, without the requirement for either aeration or strictly anaerobic conditions. It is tolerant to low pH, ethanol and salt, and has a wide growth temperature range. Moreover, it is genetically accessible, enabling metabolic engineering for cell factory optimization (Bosma, Forster, & Nielsen, 2017). The species is known to produce 1,3-propanediol, reuterin, and other related industrially important compounds in high yields from glycerol (Dishisha et al., 2014), of which reuterin has also since long been known as antimicrobial (Talarico & Dobrogosz, 1989). *L. reuteri* also has most of the genes encoding for the enzymes needed for biosynthesis of 1,2-propanediol and 1-propanol, both of which are industrially relevant chemicals. These compounds are, however, not produced under normal conditions by *L. reuteri*, requiring improved engineering- and optimization strategies to achieve commercial level cell factories and production processes (International Publication Number WO 2014/102180 AI, 2014).

Genome-scale metabolic models are highly useful for directing metabolic engineering strategies, as well as to improve understanding of the physiology and metabolism of the target organism (Rau & Zeidan, 2018; Saulnier et al., 2011). So far, highly curated and experimentally validated metabolic models have been primarily developed for model organisms such as *Escherichia coli* and *Saccharomyces cerevisiae*, but models for several LAB species are also available, including *Lactobacillus plantarum* (Teusink et al., 2006), *Lactococcus lactis* (Oliveira, Nielsen, & Förster, 2005) and *Streptococcus thermophilus* (Pastink et al., 2009). These LAB are homofermentative or facultatively heterofermentative organisms and have significant differences in metabolism compared to strict heterofermenters such as *L. reuteri*. A metabolic model for the heterofermenter *Leuconostoc mesenteroides* (Koduru et al., 2017) is available, but as it is distantly related to *L. reuteri* it is of limited use here. Models for two probiotic strains of *L. reuteri* have been previously published (Saulnier et al., 2011). They were automatically reconstructed from the same draft model we started with here (Santos, 2008). The two previously published *L. reuteri* models were used along with transcriptomics data to identify qualitative metabolic differences between the two strains as well as to analyze their probiotic properties (Saulnier et al., 2011). However, these previous models were not manually curated and were not used to quantitatively predict metabolic behavior. The construction of a genome-scale metabolic model that can be reliably used in basic research and cell factory design is a time-consuming process, requiring significant amount of manual curation and availability of strain-specific phenotypic data. At present, models obtained using automated tools or models that do not include experimental data are generally of limited use for quantitative predictions.

Here, we set out to reconstruct the metabolic network of *L. reuteri* JCM 1112, specifically for use in metabolic engineering applications, which requires collection of phenotypic data under several different conditions. We first performed an in-depth analysis of the genome to evaluate conflicting reports about metabolic pathways compared to strain DSM 20016. We then performed experiments to collect phenotypic data for the wild-type strain as well as for an alcohol dehydrogenase (*adhE*) knockout strain to constrain, validate, and test the model. The model as well as the experimental data are available in supplementary files.

## 2. Materials and methods

### 2.1 Strains, media and culture conditions

Strains used in this study are listed in Table 1 and an overview of the experimental datasets in Table 2.

**Table 1.**
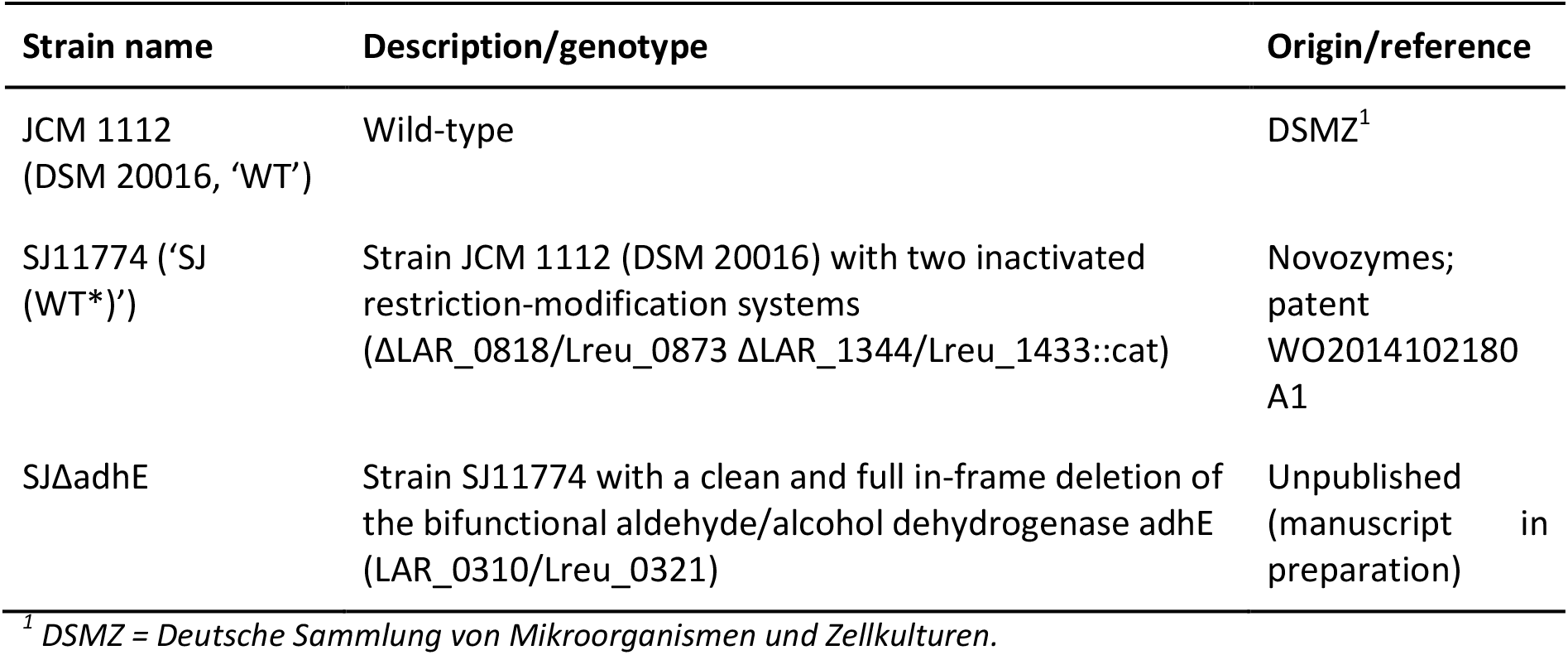
Lactobacillus reuteri strains used in this study.

**Table 2.** Experimental datasets used for the model reconstruction.

| Strain | Substrate | Growth mode | Nr of replicates | Used in: |
| --- | --- | --- | --- | --- |
| WT | Glucose | Flask | 2 | Determining energy requirements (section 2.4 and 3.1.2) |
| WT | Glucose | Reactor | 3 | Model validation (A in Figure 3) |
| WT | Glucose + glycerol | Reactor | 2 | Model validation (B in Figure 3) |
| SJ (WT*) | Glucose | Flask | 3 | Model validation (C in Figure 3) and model predictions (Figure 4a) |
| SJ (WT*) | Glucose + glycerol | Flask | 3 | Model validation (D in Figure 3) and model predictions (Figure 4b) |
| SJΔadhE | Glucose | Flask | 3 | Model validation (E in Figure 3) and model predictions (Figure 4c) |
| SJΔadhE | Glucose + glycerol | Flask | 3 | Model validation (F in Figure 3) and model predictions (Figure 4d) |

Growth curves and uptake and secretion data can be found in Additional file 1.

De Mann Rosa Sharp (MRS) medium (incl. 20 g/L glucose) was obtained from VWR and prepared according to the manufacturer’s instructions.

Chemically Defined Medium (CDM) was used as described in (Santos, 2008) / (Teusink et al., 2005) with the following modifications: arginine 5 g/L, tween-80 1 mL/L. Substrates were 111 mM glucose and 20 mM glycerol as indicated. The CDM was filter-sterilized and the final pH after mixing all components was 5.6.

All flask cultivations were performed in a stationary incubator at 37°C. A 5 mm inoculation loop of culture was inoculated from −80°C glycerol stocks into 1 mL MRS with or without glycerol in a 1.5 mL Eppendorf tube and grown overnight (16h). Next morning, cultures were washed 3x with sterile 0.9% NaCl, after which OD_600_ was measured and cells were transferred to 12 mL CDM with or without glycerol in a 15 mL Falcon tube to a starting OD_600_ of 0.08. After 4h of growth, OD_600_ was measured and cultures were transferred to a starting OD_600_ of 0.05 in 100 mL pre-warmed CDM with or without glycerol in a 100 mL Schott flask. Samples for OD_600_ measurement and HPLC were taken directly after inoculation (t=0h) and at 2, 3, 4, 5, and 6h; cultures were swirled for mixing prior to taking samples. The 6h samples were also used for protein and amino acid determinations. The time points used were all during exponential growth, ensuring a pseudo steady state (Additional file 1).

All bioreactor cultivations were performed in batch mode and samples were taken during exponential/pseudo-steady state (Additional file 1). One of the fermentations was performed in CDM at 37°C in 3.0 L bioreactors (BioFlo 115, New Brunswick Scientific/Eppendorf) with a 2.2 L working volume, 50 rpm agitation without gas sparging. The pH was controlled at 5.7±0.1 using 5N NaOH. Pre-cultures were performed similarly as for the flask cultures described above, with the pre-culture in CDM in 100 mL medium in 100 mL flasks, and reactors inoculated to an OD_600_ of 0.1. The other two reactor cultivations were performed in CDM, with and without glycerol, at 37°C in 0.4 L reactors with a 0.5 L working volume, 50 rpm agitation and sparged with N_2_ at 15 mL/min for 1h prior to inoculation. The pH was controlled at 5.8 using 5M NaOH. Fermenters were inoculated to an initial OD_600_ of 0.05 from an exponentially growing culture on CDM without glycerol. As can be seen in Additional file 1, there is no difference between the cultures in the reactors that were sparged with N2 prior to fermentations and those that were not and hence we decided to treat these as replicates.

The correlation factor between cell dry weight (gDW) and OD_600_ was experimentally determined to be 0.4007 gDW/OD_600_ in CDM and used for calculating gDW from OD_600_ in all experiments.

### 2.2 Analytical methods

Protein concentration of the cells was determined in the 6h samples as described above, via a BCA protein assay (Merck-Millipore cat. 71285) according to the manufacturer’s protocol. Prior to the BCA assay, cell pellets were washed once in 0.9% NaCl and resuspended in 0.25 mM Tris-HCl pH 7.5 and sonicated on ice with an Ultrasonic Homogenizer 300VT (BioLogics) for 3x 30s at 40% power, with 30s breaks on ice.

Amino acid composition of the cells was determined by Ansynth BV (The Netherlands) on washed cell pellets of a 6h CDM culture as described above.

Substrates, products and amino acids secreted and taken up during the cultivations were quantified using HPLC. Glucose, glycerol, ethanol, lactate, acetate, citrate, 1,2-propanediol, 1,3-propanediol, 1-propanol, 2-propanol, pyruvate, succinate and malate were quantified with either one of two HPLCs: 1) a Dionex Ultimate 3000 (Thermo Scientific) containing an LPG-3400SD pump, a WPS-3000 autosampler, a UV-visible (UV-Vis) DAD-3000 detector, and an RI-101 refraction index detector. Injection volume was 20 µL. An Aminex HPx87 ion exclusion 125-0140 column was used with a mobile phase of 5 mM H_2_SO_4_, a flow rate of 0.6 mL/min and an oven temperature of 60°C; 2) a Shimadzu LC-20AD equipped with refractive index and UV (210 nm) detectors, with an injection volume of 20 µL. A Shodex SH1011 8.0mmIDx300mm column was used with a mobile phase of 5 mM H_2_SO_4_, a flow rate of 0.6 mL/min and an oven temperature of 50°C. All amino acids, ornithine and GABA were quantified using a Dionex Ultimate 3000 (Thermo Scientific), for which the procedure is as follows: 20 µg/mL 2-aminobutanoic acid and sarcosine were used as internal standards for dilution of the samples; derivatization was performed in the autosampler. 0.5 µL sample was added into 2.5 µL of (v/v) 3-mercaptopropionic acid in borate buffer (0.4 M, pH 10.2), mixed and incubated for 20 s at 4°C to reduce free cystines. Then 1 µL of 120 mM iodoacetic acid in 140 mM NaOH was added, mixed and incubated for 20 s at 4°C to alkylate reduced cysteines. 1.5 µL of OPA reagent (10 mg o-pthalaldehyde/mL in 3-mercaptopropionic acid) was then added to derivatize primary amino acids. The reaction was mixed and incubated for 20s at 4°C. 1 µL of FMOC reagent (2.5 mg 9-fluorenylmethyl chloroformate/mL in acetonitrile) was added, mixed and incubated for 20 s at 4°C to derivatize other amino acids. 50 µL of Buffer A (Buffer A: 40 mM Na_2_HPO_4_, 0.02% NaN_3_ (w/v) at pH 7.8) at pH 7 was added to lower the pH of the reaction prior to injecting the 56.5 µL reaction onto a Gemini C18 column (3 um, 4.6 x 150 mm, Phenomenex PN: 00F-4439-E0) with a guard column (SecurityGuard Gemini C18, Phenomenex PN: AJO-7597). The column temperature was kept at 37°C in a thermostatic column compartment. The mobile phase had the following composition: Buffer A: see above, pH 7.8; Buffer B: 45% (v/v) acetonitrile, 45% (v/v) methanol and 10% (v/v) water; flow rate 1 mL/min. Derivatized amino acids were monitored using a fluorescence detector. OPA-derivatized amino acids were detected at 340_ex_ and 450_em_ nm and FMOC-derivatized amino acids at 266_ex_ and 305_em_ nm. Quantifications were based on standard curves derived from dilutions of a mixed amino acid standard (250 µg/mL). The upper and lower limits of quantification were 100 and 0.5 µg/mL, respectively.

### 2.3 Genome sequencing and analysis

For genomic DNA (gDNA) isolation, overnight cultures of DSM 20016 and SJ 11774 were grown in MRS and the pellet was used for gDNA isolation using the Epicentre MasterPure^TM^ Gram Positive DNA Purification kit according to the manufacturer’s protocol. Subsequent genome sequencing was performed at the sequencing facility at the NNF Center for Biosustainability. Library preparation was performed using KAPA HyperPlus Library Prep Kit (ROCHE) with Illumina-compatible dual-indexed PentAdapters (PentaBase). The average size of the library pool was 317 bp. Sequencing was performed on MiSeq (Illumina) using the MiSeq Reagent Kit v2, 300 Cycles (Illumina). The libraries were loaded to the flow cell at 10 pM and sequenced using paired-end reads of 150 bp. Read quality check was performed with FastQC version 0.11.5. Mutations relative to reference (*L. reuteri* JCM 1112, GenBank accession nr AP007281, annotated with Prokka version 1.11) were identified using Breseq (version 0.31.0) (Deatherage & Barrick, 2014). Mean coverage was 143.7x (SJ 11774) and 129.5x (DSM 20016). All runs were performed at Danish national supercomputer for life sciences (Computerome), Technical University of Denmark. For this work, the annotated genome of *L. reuteri* JCM 1112 from NCBI was used. During the reconstruction, several genes were re-annotated, based on BLAST and physiological data. A list of all genes in the JCM 1112 genome can be found in Additional fie 2, along with annotations from the GenBank file and which model reactions are associated with each gene.

### 2.4 Metabolic reconstruction

The *L. reuteri* JCM 1112 metabolic reconstruction was based on an unpublished, automatically generated draft reconstruction of JCM 1112 (Santos, 2008). We performed extensive manual curation, including: gap filling, updating and adding gene-protein-reaction (GPR) associations, updating gene IDs, updating metabolite- and reaction abbreviations, in line with the BiGG database (King et al., 2016), updating and adding missing formulas and/or charges to metabolites, fixing unbalanced reactions, adding annotation to metabolites, reactions and genes and detailed review and integration of organism specific data. A biomass objective function was formulated based on available data on *L. reuteri* and related strains. The ATP cost of growth-associated maintenance (GAM) was estimated using one of the data sets (Table 2) by adjusting the GAM parameter so that growth predictions matched *in vivo* growth. This data set was then excluded from subsequent validation and prediction steps.

### 2.5 Flux balance analysis

Flux balance analysis (FBA) was used to analyze the genome-scale metabolic model (Fell & Small, 1986; Savinell & Palsson, 1992) by constraining exchange reactions in the model with experimental values of substrate uptake and secretion rates. To take into account that the Embden–Meyerhof– Parnas (EMPP) pathway is a minor glycolytic pathway in *L. reuteri* compared to the phosphoketolase pathway (PKP) (section 3.1.1), an additional flux constraint was added to the model

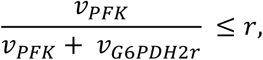

where *r* is an empirically determined flux ratio, **v**_PFK_ denotes flux in the rate limiting step of the EMPP and **v**_G6PDH2r_ is the flux in the first reaction branching into the PKP.

We used a variant of FBA called parsimonious FBA (Lewis et al., 2010) which identifies flux values corresponding to maximum growth with the side constraint that the sum of absolute flux values is made as small as possible. The sum of fluxes is proxy for enzyme usage and the method can therefore be considered to simulate biological pressure for rapid and efficient growth using minimum amount of resources (enzymes). An advantage over FBA is that the resulting solution is likely to contain fewer infeasible flux cycles. Model simulations were carried out in Python with the CobraPy toolbox (Ebrahim, Lerman, Palsson, & Hyduke, 2013) and GLPK solver. All code used in the simulations is provided in the form of a Jupyter notebook in Additional file 3 and on https://github.com/steinng/reuteri. The Escher package (King et al., 2015) was used for visualization of flux predictions. Escher maps of *L. reuteri*’s central metabolism are provided in Additional file 4, both simplified maps as shown in sections 3.2.2 and 3.2.3 as well as a detailed map linking different sugar utilization pathways to the central metabolism.

To predict growth rates the model was constrained with uptake rates of glucose, glycerol and five amino acids (Arg, Ser, Asn, Asp and Glu), and with the secretion rates of ethanol, lactate, acetate and 1,3-propanediol. Effects of knocking out the *adhE* gene were predicted by temporarily deleting it from the network. Where the effects of an active 1,2-propanediol pathway were predicted, a methylglyoxal synthase (MGS) was added to the model and optimized for growth.

To predict the theoretical maximum yields of selected target compounds, a reaction enabling the secretion of the corresponding metabolite was added to the model, unless an exchange reaction already existed, and flux through the reaction maximized. The glucose uptake rate was 25.2 mmol gDW^−1^ h^−1^, based on experimental data, and free secretion of by-products was allowed. For the production of L-alanine, an L-alanine dehydrogenase was added to the model. The production of ethyl lactate required the addition of a lactate acyl transferase and a reaction for the condensation of lactoyl-CoA with ethanol (Lee & Trinh, 2018). To produce 1-propanol, a methylglyoxal synthase (MGS) was added to the model. The presence of a complete 1-propanol pathway enables more efficient regeneration of NAD and the flux predictions were therefore repeated in the presence of an active MGS. To simulate a non-limiting phosphofructokinase, the flux constraint involving **v**_PFK_ above was omitted.

## 3. Results and discussion

### 3.1 Metabolic network reconstruction

To reconstruct a genome-scale metabolic model of *L. reuteri* suitable for use in cell factory design and optimization, we built upon a draft metabolic model of *L. reuteri* JCM 1112 described in (Santos, 2008) that we in turn extensively curated. The Memote tool (Lieven et al., 2018) was used to assess the quality of the reconstruction and to guide the curation process (Additional file 5). The main characteristics of the resulting Lreuteri_530 model (Additional file 6) are listed in Table 3.

**Table 3.** Main characteristics of Lreuteri_530 - the L. reuteri JCM 1112 genome-scale metabolic reconstruction.

| Genome characteristics |  |
| --- | --- |
| Genome size | 2.04 Mb |
| Total protein coding sequences | 1943 |
| Model characteristics |  |
| Genes | 530 |
| Percentage of genome | 27% |
| Reactions (with GPR) | 710 (690) |
| Metabolites (unique) | 658 (551) |
| Memote total score | 62% |

#### 3.1.1 Curation process

Reactions and metabolites were abbreviated according to the BiGG database nomenclature where applicable and annotations with links to external databases included. Genes from the JCM 1112 genome were identified with locus tags from the GenBank file, and annotations were included which contain: the old locus tag which is often found in older literature, the NCBI protein ID, gene annotation and the protein sequence. Apart from general network curation, organism-specific information obtained from laboratory experiments and from available literature was integrated by reviewing reactions, genes and gene-protein-reaction (GPR) rules.

##### Resequencing reveals inconsistencies between the “same” strains *L. reuteri* DSM 20016 and JCM 1112 - implications for glycolytic genes

The two most well-known strain names and origins for the type strain are DSM 20016 and JCM 1112 from the DSMZ and JCM culture collections, respectively. These two are derived from the same original human faeces isolate *L. reuteri* F275 (Kandler, Stetter, & Köhl, 1980), which was grown and stocked in two different laboratories (Frese et al., 2011). Both genomes have been sequenced previously and a comparison showed two remarkable differences between these two strains derived from the same parent strain: DSM 20016 was missing two large regions (Morita et al., 2008), most likely lost during the 20 years of separate laboratory cultivation (Frese et al., 2011). The first region (8,435 bp, flanked by IS4 insertion sequences on each end) contains genes for glycolysis, namely glyceraldehyde-3-P dehydrogenase, phosphoglycerate kinase, triosephosphate isomerase, and enolase. The second region (30,237 bp, flanked by two different insertion sequence elements) contains a gene cluster for nitrate reductases and molybdopterin biosynthesis (Morita et al., 2008). As the first island consists of glycolytic genes, the implications of its presence or absence are profound. This island is absent in DSM 20016, but we could identify homologs of all this island’s genes except glyceraldehyde-3-P dehydrogenase elsewhere in its genome based on annotation and/or BLAST.

During the preparation of our model, it became clear that there are inconsistencies in naming and hence gene content of the *L. reuteri* type strain. We sequenced the DSM 20016 strain that we obtained from DSMZ and this showed that its genome is identical to that of JCM 1112 instead. A similar result of these strains being ‘swapped’ was obtained by others based on whole genome sequencing (US 2015O125959A1, 2015) and PCR of part of the largest missing region in DSM 20016 (Etzold et al., 2014). This inconsistency between the two strains does not seem to be commonly known and taken into account, and we suspect that some papers referring to either the DSM or the JCM strain might in fact be working with the other strain. For example, the DSM 20016 strain used by Sun et al., sequenced in 2015 (accession nr AZDD00000000), contains the islands and hence is actually the JCM 1112 strain (Sun et al., 2015), whereas the DSM 20016^T^ referred to by Morita et al., sequenced in 2007 by JGI (accession nr CP000705), was shown to be DSM 20016, missing the islands (Morita et al., 2008). Both strains were obtained from DSMZ. This highlights the importance of re-sequencing of strains ordered from culture collections or lab strains present in the laboratory before using them for engineering or characterization studies. We strongly suggest that studies working with any *L. reuteri* type strain perform PCR on the two islands or perform resequencing as the presence of the first island determines whether the strain contains a full glycolytic pathway or not. In our model, we have included all genes in the islands based on the sequencing results. The reconstruction was based on the genomic information of the JCM 1112 strain, obtained from NCBI, and the genes in the model are identified with the locus tags obtained from there. As many other publications refer to genes in the DSM 20016 strain or use the old locus tags from the JCM 1112 genome, we have included a table (Additional file 2) which lists: the locus tags used in the model (LAR_RSXXXXX), the old locus tags (LAR_XXXX), the annotations obtained from the NCBI GenBank file, the NCBI protein IDs (WP numbers), the locus tags of the corresponding genes in the DSM 20016 strain, when applicable (Lreu_XXXX), and finally the reaction(s) in the metabolic model associated with the genes.

##### Phosphofructokinase (PFK) and the distribution between EMP and PK pathway usage

Obligately heterofermentative lactobacilli like *L. reuteri* are often considered to solely use the phosphoketolase pathway (PKP) instead of the Embden-Meyerhof-Parnas pathway (EMPP) for glucose consumption (Bosma et al., 2017) (Figure 1). Both pathways result in the glycolytic intermediate glyceraldehyde-3-phosphate but use different redox cofactors (Figure 1). As the PKP yields one and the EMPP two molecules of glyceraldehyde-3-phosphate, the PKP has a lower energy yield than the EMPP (Figure 1). The PKP generally results in the production of one molecule of lactate and one molecule of ethanol or acetate for one glucose molecule while the EMPP generally yields two lactate molecules. Key enzymes of the EMPP are fructokinase (FK), glucose-6-phosphate isomerase (PGI), phosphofructokinase (PFK), fructose-bis-phosphate aldolase (FBA), and triosephosphate isomerase (TPI). In line with the idea that heterofermenters use the PKP, Sun et al. showed in a comparison of 213 LAB genomes that *pfk* was lacking from a distinct monophyletic group formed by mainly (87%) obligatively and otherwise facultatively heterofermentative *Lactobacillus* spp., including *L. reuteri* DSM 20016 and *L. panis* DSM 6035 (Sun et al., 2015). Contrary to most other species in the same group, these two species did contain *fba*, which has traditionally been linked to the presence of the EMPP. Despite the absence of *pfk*, EMPP activity has been observed in several *L. reuteri* strains and in some strains it appears to play a major role compared to the PKP, depending on the growth phase, and showing strain-specific differences (Årsköld et al., 2008; Burgé et al., 2015). For modeling and engineering purposes, it is crucial to understand the presence and activity of the PKP vs the EMPP.

**Figure 1.**
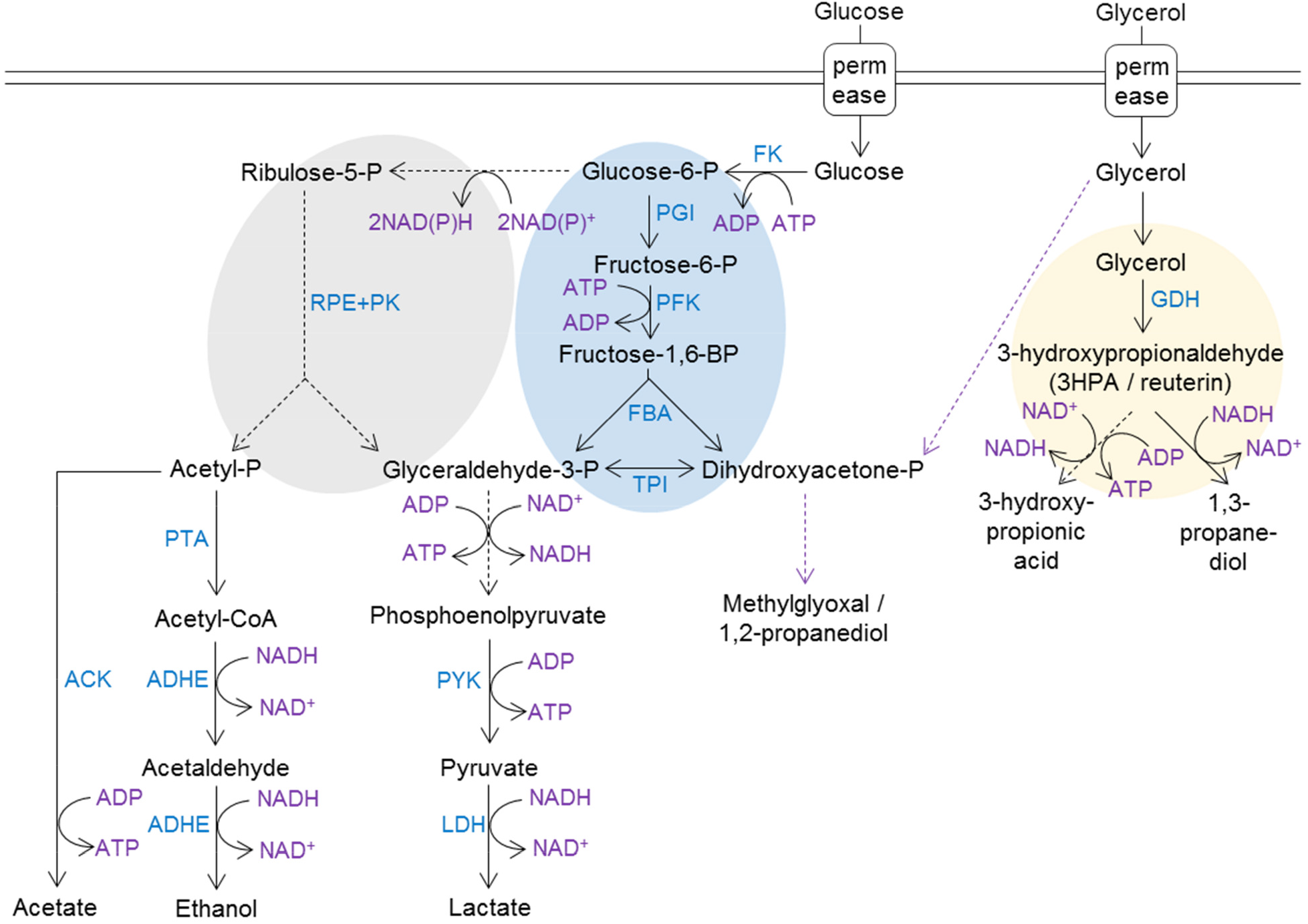
Condensed overview of the central metabolism in L. reuteri. Dotted purple arrows indicate pathways for which genes or homologs are present but likely not active in L. reuteri JCM 1112. Dotted black arrows indicate multiple enzymatic steps. Yellow background circle indicates microcompartment; blue background indicates the EMP pathway; grey background indicates the phosphoketolase pathway. Abbreviations: FK: fructokinase/glucokinase; PGI: glucose-6-phosphate isomerase; PFK: phosphofructokinase; FBA: fructose-bis-phosphate aldolase; TPI: triosephosphate isomerase; PGM: phosphoglucomutase; SP: sucrose phosphorylase; M2DH: mannitol-2-dehydrogenase; RPE+PK: ribulose epimerase + phosphoketolase; GDH: glycerol dehydratase I. Adapted from (Bosma et al., 2017).

Årsköld et al. (2008) compared the genomic organization of 13 sequenced *Lactobacillales* and showed that *L. reuteri* (strains ATCC 55730 and DSM 20016) is one of the four exceptions that do not have a *pfkA* gene where this is located in all other species. Nevertheless, they detect PFK and EMPP activity in strain ATCC 55730 and subsequently identify two genes (GenBank accession nrs EF547651 and EF547653) for orthologues of *pfkB*, a minor PFK-variant in *E. coli* (Årsköld et al., 2008). In analogy with Årsköld et al. in *L. reuteri*, Kang et al. (Kang, Korber, & Tanaka, 2013) identified a ribokinase in the obligately heterofermentative *L. panis* PM1 with 82% similarity to the *pfkB* gene identified in *L. reuteri* ATCC 55730 from Årsköld et al. (74% in our own BLAST search).

A BLAST comparison of the *pfkB* protein sequence of *L. panis* PM1 (GenBank accession nr AGU90228.1) and *L. reuteri* ATCC 55730 (GenBank accession nr ABQ23677.1) against *L. reuteri* JCM 1112 resulted in 81% and 99% identity, respectively, to JCM 1112 gene number LAR_RS02150, which is annotated as ribokinase rbsK_2. On a gene level, this gene shares 97% identity with *L. reuteri* ATCC 55730 and 73% with *L. panis* PM1. The same identities were found in *L. reuteri* DSM 20016 for gene LREU_RS02105 (previously Lreu_0404, GenBank protein KRK49592.1). A second gene annotated as “ribokinase rbsK_3” (locus tag LAR_RS06895) showed only limited query coverage and identity and hence rbsK_2 is the most likely homolog of *pfkB*. The growth experiments conducted in the present study with JCM 1112 are in line with the findings of Burgé et al. and indicate minor though detectable usage of the EMPP in this strain with a peak in the early growth stage (Figure 2), in which this rbsK_2 likely fulfills the role of *pfkB*. The average flux through the EMPP in all cultures was 7.0% (Figure 2) and was used to define the corresponding flux split ratio in the model (section 2.5).

**Figure 2.**
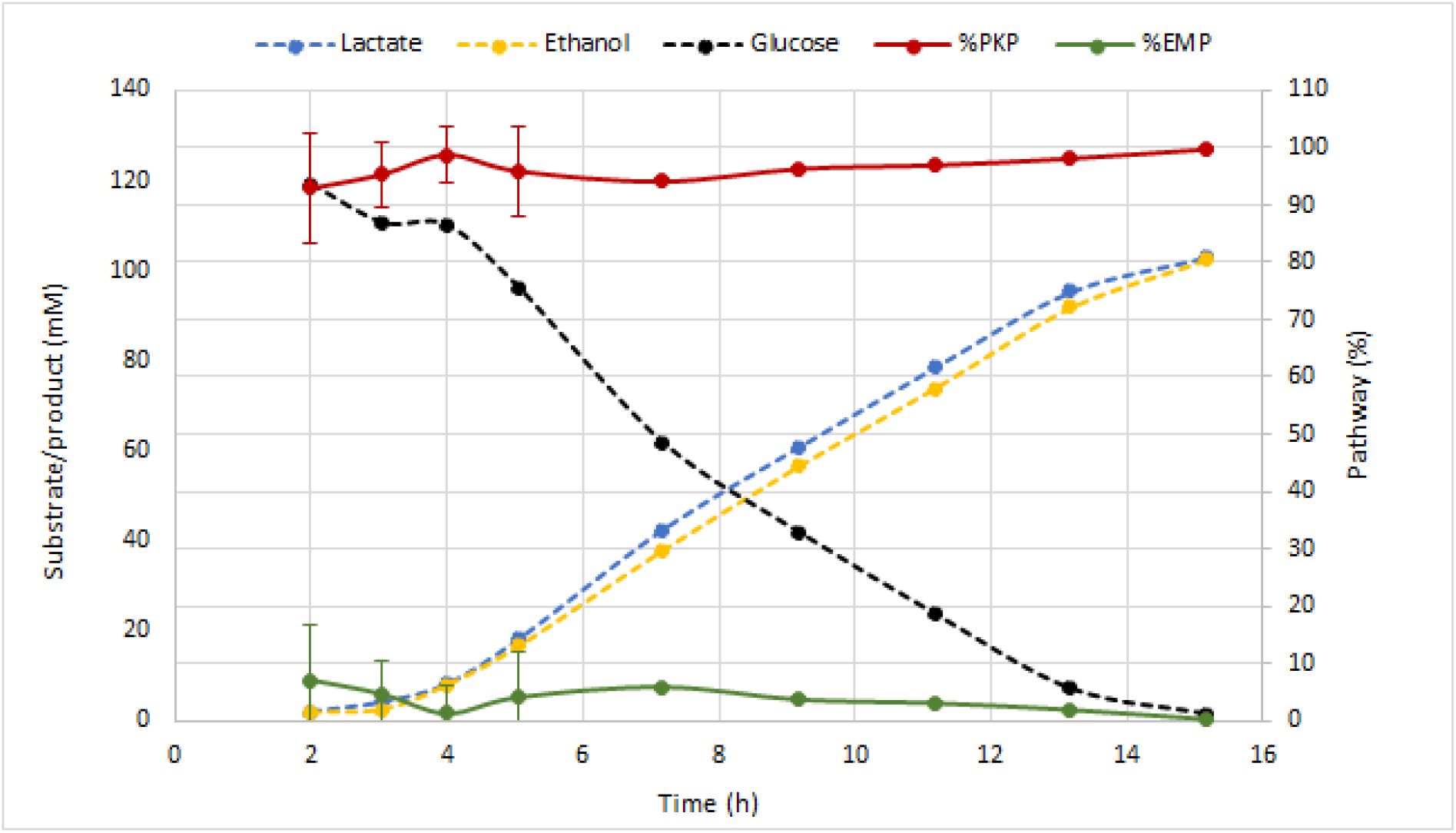
Typical fermentation profile and distribution between the EMP and PK pathways in L. reuteri JCM 1112 in chemically defined medium with glucose as the sole carbon source. Data are averages of the all the datasets used to constrain and validate the model, with error bars representing standard deviation. The percentage of PKP usage was defined as in Burgé et al., i.e. as the ethanol concentration divided by the sum of lactate and ethanol concentrations divided by 2.

##### Sugar transport

Transport of carbohydrates can be mediated by ATP-Binding Cassette (ABC) transporters, phosphotransferase systems (PTS), or secondary transporters (permeases of the Major Facilitator Superfamily, MFS) (Saier, 2000). PTS systems mediate hexose mono- or dimer transport and phosphorylation simultaneously – mostly by using PEP to pyruvate conversion as phosphate donor, whereas ABC-transporters (mostly used for pentoses) and permeases (both pentoses and hexoses) perform only transport, and a separate ATP-utilizing kinase step is needed for sugar phosphorylation. Moreover, in Gram positives, PTS systems have an important role in carbon catabolite repression via phosphorylation cascades and direct interaction with the carbon catabolite repression protein A (ccpA) (Galinier & Deutscher, 2017; Görke & Stülke, 2008). Heterofermentative LAB contain fewer PTS system components than homofermentative LAB, which is thought to be the result of gene loss (Zheng, Ruan, Sun, & Gänzle, 2015). In general, organisms using the EMPP are believed to use PTS systems, and organism using the PKP to use secondary carriers (Romano, Trifone, & Brustolon, 1979). Likely as a result of the lack of full PTS systems, glucose utilization is not constitutive but substrate-induced in heterofermenters, and utilization of several other sugars is not repressed by glucose (Galinier & Deutscher, 2017). Sugar transport in heterofermenters is poorly characterized, and only recently a study was dedicated to the genomic and phenotypic characterization of carbohydrate transport and metabolism in *L. reuteri*, as representative of heterofermentative LAB (Zhao & Gänzle, 2018). This showed that *L. reuteri* completely lacks PTS systems and ABC-transporters and solely relies on secondary transporters of the MFS superfamily, which use the proton motive force (PMF) as energy source for transport (Zhao & Gänzle, 2018). In *L. reuteri* JCM 1112, we could identify the two common proteins of the PTS system, Enzyme I (Lreu_1324) and HPr (Lreu_1325). Some sugar-specific parts were present, but no complete PTS was identified. As a result, all sugar transport in the model takes place via the PMF.

##### Glycerol utilization

*L. reuteri*, like many lactobacilli, is known to be unable to grow on glycerol as a sole carbon source, but can use it as an alternative electron acceptor, providing a means to gain energy on a variety of carbon sources (Sriramulu et al., 2008; Talarico, Axelsson, Novotny, Fiuzat, & Dobrogosz, 1990). *L. reuteri* is the only known lactobacillus producing large amounts of 3-hydroxypropionaldehyde (reuterin, 3-HPA) from glycerol. This is an intermediate in the pathway to 1,3-propanediol (1,3-PDO, also produced by *L. reuteri*, depending on the conditions used) that is known to be toxic and produced in a microcompartment (Chen, Bromberger, Nieuwenhuiys, & Hatti-Kaul, 2016). The reason why it cannot grow on glycerol as sole carbon source is currently not fully clear, although it is likely related to gene regulation. All the genes that are necessary to convert glycerol to dihydroxyacetone phosphate via either dihydroxyacetone (DHA) or glycerol-3-phosphate and hence shuttle it into glycolysis are present in the *L. reuteri* genome (Chen et al., 2016). However, several of these genes have been shown to be downregulated in the presence of glycerol (Chen et al., 2016; Santos et al., 2008). Furthermore, the *L. reuteri* glycerol dehydrogenase also has activity as 1,3-PDO:NAD-oxidoreductase, whereas in for example *Klebsiella pneumoniae*, which does produce glycolytic end products from glycerol, these are two different enzymes (Talarico et al., 1990). It seems that the physiological role of this enzyme in *L. reuteri* is the reduction of 3-HPA to 1,3-PDO, rather than glycerol to DHA conversion, explaining the lack of growth on glycerol (Talarico et al., 1990).

##### Other pathways

Most heterofermentative LAB possess a malolactic enzyme but no malic enzymes (Landete, Ferrer, Monedero, & Zúñiga, 2013), which is also the case for our *L. reuteri* strain, based on sequence comparisons with the *L. casei* strain used by Landete et al. (Landete et al., 2013). Based on BLAST analysis and in line with literature, *L. reuteri* JCM 1112 possesses a malate dehydrogenase and PEP carboxykinase, and cannot utilize citrate; malate (and fumarate) is converted to succinate (Gänzle, Vermeulen, & Vogel, 2007).

From a biotechnological perspective, an interesting branch point of central carbon metabolism is the conversion from methylglyoxal (MG) to 1,2-propanediol (1,2-PDO), which can then be further metabolized into 1-propanol and propanoate. *L. reuteri* possesses all enzymes needed for these pathways, except methylglyoxal synthase (MGS), the step of the pathway, converting dihydroxyacetone phosphate into MG (Gandhi, Cobra, Steele, Markley, & Rankin, 2018; Sriramulu et al., 2008). It has been shown that when MG is added to *L. reuteri* JCM 1112 cultures or when a heterologous *mgs* is expressed, all the subsequent metabolites are formed (International Publication Number WO 2014/102180 AI, 2014). Although we identified a potential distant homolog of *mgs* in the *L. reuteri* genome, this homolog is clearly not active under normal conditions since no 1,2-PDO was observed in our experiments. Hence, all the genes in these pathways except *mgs* were included in the reconstruction. For methylglyoxal reductase, *mgr*, we also identified several aldo/keto reductases as possible homologs, based BLAST comparison to genes identified in (Gandhi et al., 2018). However, verification of these hypothetical activities would need extensive enzyme assays, and it is also likely that this reaction is performed by LAR_RS09730 (Glycerol dehydrogenase) (Altaras & Cameron, 1999; Yamada & Tani, 2011), this has been added to the reconstruction for the MGR reaction. Alternatively, MG might be converted directly to lactate by a glyoxalase (Gandhi et al., 2018).

*L. reuteri* can produce vitamin B12, and the structure and biosynthetic genes have been studied (Santos et al., 2007, 2008). The corresponding pathway is present in the reconstruction and is active during growth predictions.

#### 3.1.2 Biomass reaction and energy requirements

A biomass objective function (BOF), which contains all necessary components for biomass biosynthesis, is commonly used to predict growth rate in metabolic models. Ideally, the BOF should be constructed based on organism-specific experimental data, mainly the fractional composition of the macromolecules (proteins, DNA, RNA, lipids, etc.) and their individual building blocks (amino acids, nucleotides, fatty acids, etc.), as well as the energy necessary for their biosynthesis (Feist & Palsson, 2010). The protein fraction is a significant fraction of the biomass and was therefore measured. The remaining macromolecular fractions were derived from *L. plantarum* (Teusink et al., 2006) and *L. lactis* (Oliveira et al., 2005). The ratio of amino acids in the *L. reuteri* biomass was also measured. Nucleotide composition was estimated from the genome, which in the case of RNA is not ideal since it assumes equal transcription of all genes. We however preferred to use this approximation instead of using experimental data from another organism. Fatty acid composition of *L. reuteri* was obtained from literature (Liu, Hou, Zhang, Zeng, & Qiao, 2014), while phospholipid composition was adopted from *L. plantarum*. The composition of lipoteichoic acid (Walter et al., 2007) and exopolysaccharides (Ksonzeková et al., 2016) in *L. reuteri* were obtained from literature. Peptidoglycan composition was adopted from *L. plantarum* and glycogen was assumed to be negligible (Dauner & Sauer, 2001; Dauner, Storni, & Sauer, 2001).

Energy required for growth-(GAM) and cell maintenance (NGAM) are important parameters in metabolic models, and can be estimated from ATP production rates, which can be calculated from experimental data obtained at different dilution rates (Tempest & Neijssel, 1984). Unfortunately, this data is not publicly available for *L. reuteri*. These parameters have been estimated from experimental data for several other LAB, including *L. plantarum*, and reported in literature (Teusink et al., 2006). Even though *L. reuteri* and *L. plantarum* are relatively closely related, adopting these parameters from *L. plantarum* can negatively affect the quality of model predictions. When the differences in physiologies of *L. plantarum* and *L. reuteri* are considered, it is possible that *L. reuteri* requires less energy: (1) The genome is only ~2 Mb, while *L. plantarum*’s genome is 3.3 Mb. (2) *L. reuteri* is an obligate heterofermenter, which means it uses almost solely the PKP (Fig. 2) to break down glucose, resulting in one ATP per glucose, while a facultative heterofermenter like *L. plantarum* uses the EMPP when grown on glucose, resulting in two ATPs. (3) LAB in general have low catabolic capabilities, and for *L. reuteri* this includes auxotrophy for several amino acids. This, combined with the fact that macromolecular biosynthesis is already accounted for in the model reactions, supports the claim that adopting energy parameters from *L. plantarum* can negatively affect model predictions, as we also observed when evaluating this in our model. We decided to use one of our experimental datasets (Table 2) to estimate the GAM value, while using the NGAM value from *L. plantarum* (section 2.4). In general, NGAM represents only a small portion of the total energy requirements of the cell and therefore has much smaller effect on model predictions than GAM. This resulted in a GAM value of 10.2 mmol gDW^−1^ h^−1^. Detailed description of the biomass reaction, relevant data and calculations can be found in Additional file 7.

### 3.2 Model applications

#### 3.2.1 Model validation using experimental data: Growth rate comparisons

To validate the model, several different datasets (Table 2) with measured uptake- and secretion rates of carbon sources, amino acids and organic byproducts were used to constrain exchange fluxes in the model. The predicted growth rates were compared with observed experimental growth rates (Figure 3). In all cases, flux through the EMPP was set to maximally 7% based on the experimentally determined value (Figure 1). The chemically defined culture medium used in the growth experiments contained all 20 amino acids, except for L-glutamine. Subsequently, all these amino acids were quantified during growth and the model was constrained with the resulting uptake rates. Of all the amino acids, only arginine was depleted at the end of the exponential phases in data sets A, B and C (Additional file 1). Due to auxotrophy for several amino acids (Glu, His, Thr, Arg, Tyr, Val, Met, Try, Phe, Leu), the model is highly sensitive to uncertainties in measurements, as well as in determined protein- and amino acid fractions of the biomass reaction. To accurately represent amino acids in the biomass reaction, both the protein content and the amino acid ratio were measured (Additional file 7). By enabling unrestricted uptake of amino acids in the model, we noticed that only 5 amino acids (Arg, Ser, Asn, Asp, Glu) needed to be constrained with measured uptake rates for accurate growth predictions, for both the wild-type and the mutant. This is due to their role in energy- and cofactor metabolism, not only in biomass biosynthesis. Hence, only this minimum number of amino acids was used to constrain the model in the following. The remainder were assumed to be non-limiting by allowing unrestricted uptake. This has twofold advantage. First, it limits the effects of uncertainties in amino acid uptake rate measurements on model predictions, a problem exacerbated by the amino acid auxotrophy. Second, it simplifies future applications of the model by reducing the number of measurements needed.

**Figure 3.**
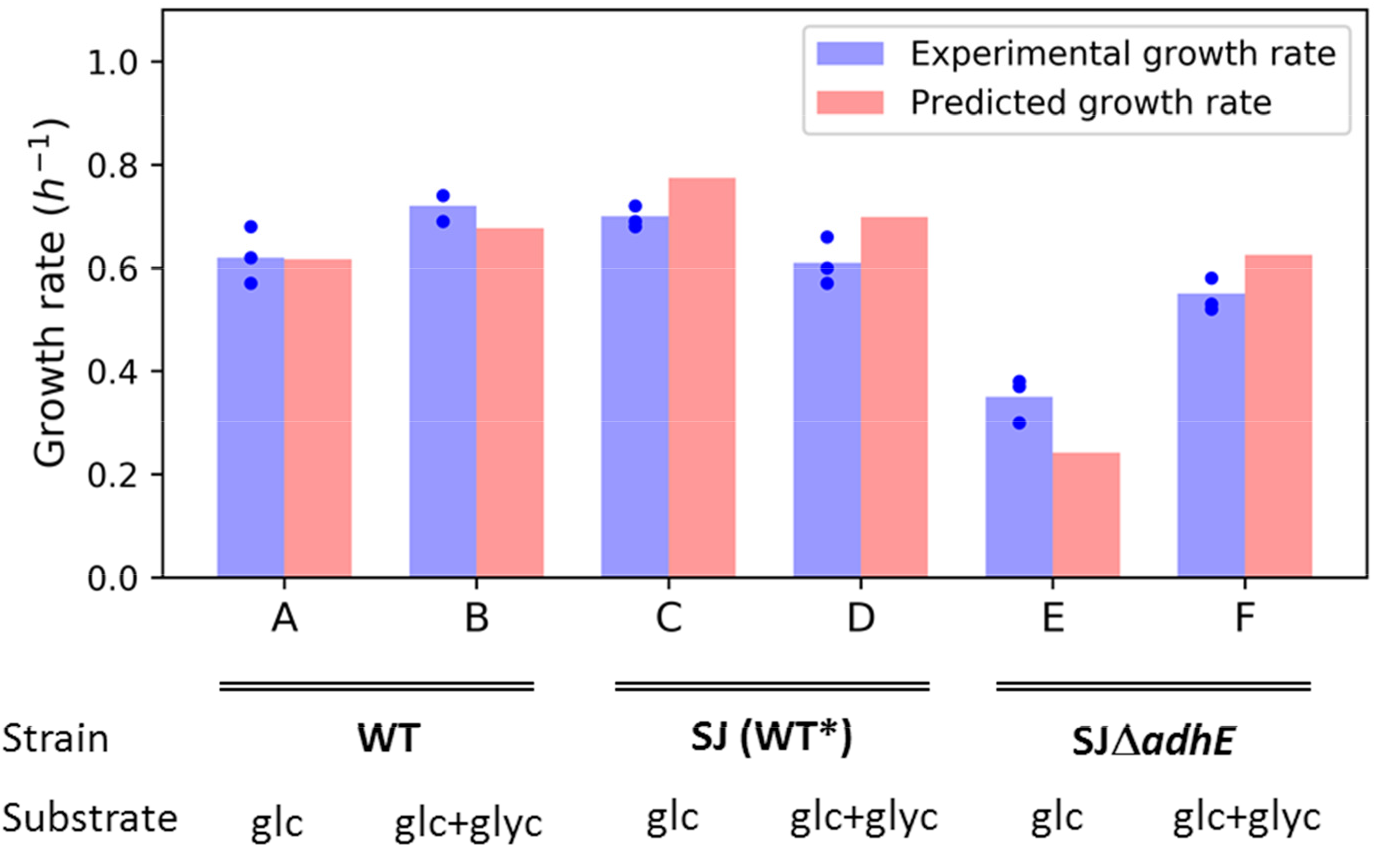
Predicted and experimental growth rates. Experimentally measured growth rates for each of the six data sets are shown in blue, with blue dots denoting individual replicates and blue bars representing average values. For each dataset, the model was constrained with average experimental values for uptake and secretion rates of carbon sources, byproducts and selected amino acids, and optimized for growth. Predicted growth rates are represented by red bars. Different datasets used are indicated with letters - abbreviations: glc: glucose; glyc: glycerol.

In most cases, model predictions and *in vivo* data were in good agreement (Figure 3). Datasets C and D in Figure 3 show a variant of the WT strain (marked SJ (WT*)), which lacks two restriction modification (RM) systems for easier genetic manipulation (Table 1). Datasets E and F show a mutant derived of the SJ strain with a clean and in-frame deletion of the *adhE* gene (bifunctional aldehyde/alcohol dehydrogenase). The model predicts slightly higher growth rates than observed *in vivo* for the SJ strain (datasets C and D in Figure 3) and the mutant strain grown on glucose and glycerol (F in Figure 3). Unexpectedly, the RM-modifications in the SJ strain seem to slightly alter its behavior on CDM with glucose and glycerol compared to the WT (Additional file 1). For the mutant strain grown on glucose (dataset E in Figure 3), the model predicts a slightly lower growth rate than observed *in vivo*, though both show a large decrease in growth, compared to the WT. The most likely explanation for this is that some glucose is being taken up *in vivo*, even though the measurements did not show this (the likely amount consumed between two samples is within the error of the assay). Secretion of 2.6 mmol gDW^−1^ h^−1^ of lactate and 2.7 mmol gDW^−1^ h^−1^ of acetate was observed *in vivo*. The model, however, does not predict lactate and acetate secretion unless some glucose uptake is allowed. If a glucose uptake of 2.6 mmol gDW^−1^ h^−1^ is allowed, the growth rate increases from 0.22 to 0.34 h^−1^, compared to 0.30 h^−1^ *in vivo.* Amino acid measurements showed that the mutant in dataset E used L-arginine to a greater extent than the WT, which the model predicts is used to generate energy via the arginine deiminase pathway, resulting in increased growth.

#### 3.2.2 Effects of adding glycerol and deleting *adhE*

To investigate the applicability of the model for cell factory design, it was used to predict the effects of adding glycerol to the glucose-based culture medium, as well as knocking out the *adhE* gene, which plays a critical role in ethanol production and redox balance (Figure 1). The datasets used here are the same as in section 3.2.1 (datasets C - F in Figure 3). There, the aim was to validate the model by means of comparing predicted growth rates to experimentally determined growth rates. In this section, we look more specifically at predicted flux distributions in central metabolism, both with and without strain- and condition-specific experimentally determined constraints. For this purpose, we studied two cases in order to answer the following questions: (1) If the model is constrained only with experimentally determined glucose- and five amino acid uptake rates from the WT strain grown on glucose, how do the predicted effects of glycerol addition and/or *adhE* knock-out (dark green bars in figure 4) compare to *in vivo* growth rate and uptake- and secretion measurements (light orange bars in figure 4)? This was tested to evaluate the applicability of the model in a practical setting. One of the main goals of using a model like this should be to probe the effects of genetic and media perturbations *in silico*, *i.e.* without having to do extensive condition-specific cultivations and measurements beforehand. (2) If the model is constrained with uptake- and secretion rates of carbon source(s), amino acids and byproducts of the strain and condition under study, how well do the model predictions (light green bars in figure 4) compare to *in vivo* results? Here the model was allowed, but not forced, to take up (lower bound constrained, upper bound unconstrained) and secrete (lower bound unconstrained, upper bound constrained) metabolites according to the experimental data. This tells us if the model, when imposed with realistic limitations, “chooses” a flux distribution which results in extracellular fluxes of metabolites in line with *in vivo* data. In both cases, the constrained amino acids only included Arg, Ser, Asn, Asp and Glu as before (see section 3.2.1) and in case 1 the allowed glycerol uptake rate was arbitrarily limited to 25 mmol gDW^−1^ h^−1^, when glycerol effects were being predicted.

**Figure 4.**
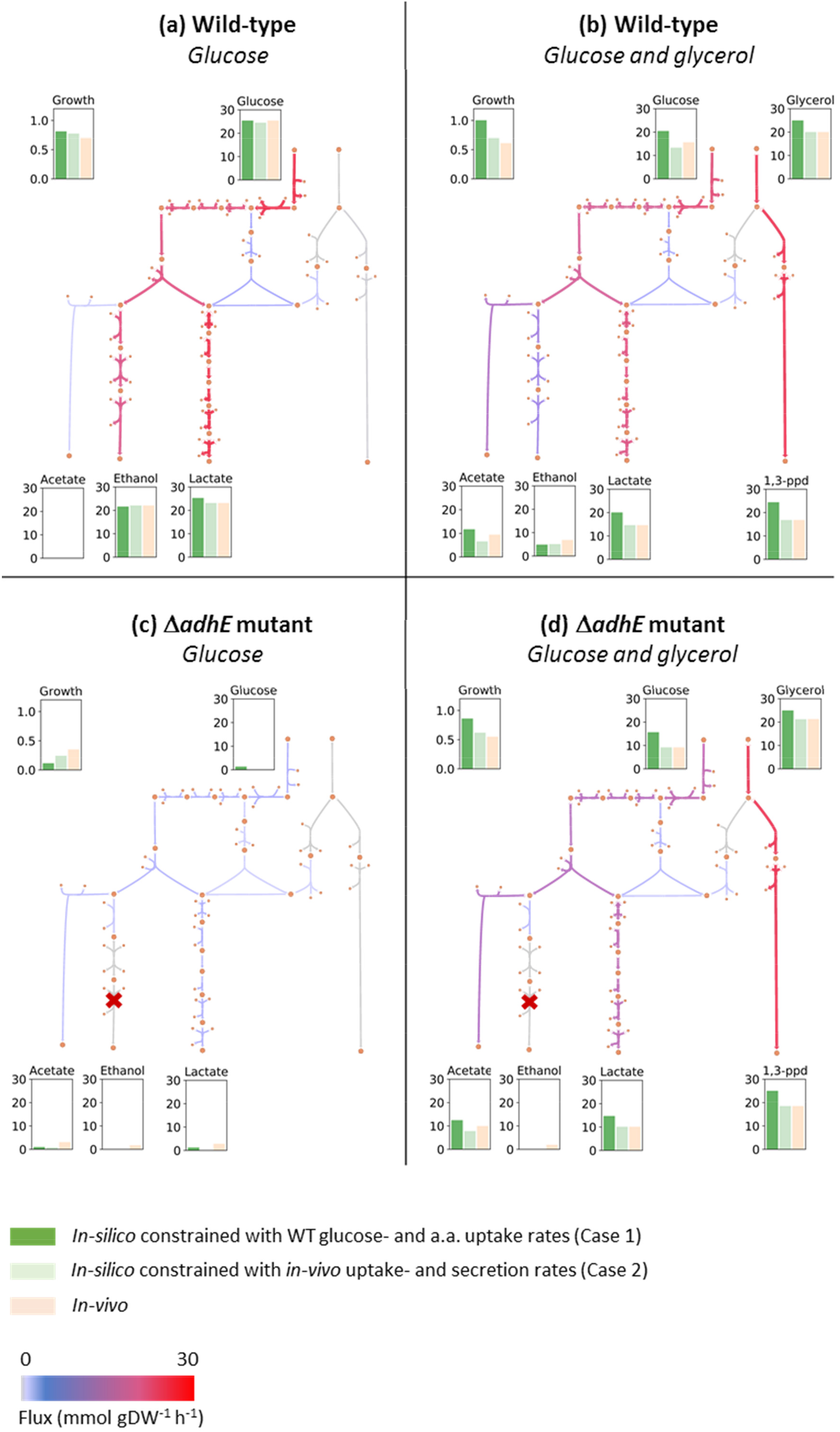
Predicted and experimental fluxes of key metabolites in the wild-type strain (SJ) and the adhE mutant. The wild-type strain was grown on glucose (a) and glucose and glycerol (b), and the adhE mutant was also grown on glucose (c) and glucose and glycerol (d). Bar plots show the average measured rates from 3 replicates (light orange), predicted rates from model constrained with average experimental uptake rates of the WT grown on glucose, or case 1 (dark green), and predicted rates from model constrained with average experimental rates from the strain and condition under study, or case 2 (light green). Metabolic maps show predicted flux distributions for case 1. All units for uptake- and secretion rates are in mmol gDW^−1^ h^−1^ and for growth rates in h^−1^.

The flux maps in Figure 4 show results for case 1 (dark green bars). The predicted uptake of glucose and glycerol (dark green bars in figure 4b) is higher than observed *in-vivo* (light orange bars in figure 4b), resulting in higher secretion of by-products and a higher growth rate as well. However, the distribution of secreted by-products is very similar. The effect of glycerol can be predicted quite well with the model as ethanol secretion decreases and acetate secretion increases, relative to glucose uptake, and 1,3-propanediol is secreted in large amounts (compared to graphs in Figure 4a). Several studies have described an increased growth rate in *L. reuteri* when glycerol is added to a glucose-based medium (in flasks and bioreactors), which is to be expected based on inspection of redox balance (Chen et al., 2016; Santos, 2008; Talarico et al., 1990) and this is also what we observed *in silico* in case 1. However, *in vivo* we consistently observed a small decrease in growth rate for this strain when glycerol was added (Additional file 1).

In line with existing literature reports (Chen et al., 2016), knocking out the *adhE* gene has dramatic effects on the metabolism when glucose is the sole carbon source, both *in vivo* and *in silico* (Figure 4c). This is due to redox imbalance since AdhE no longer recycles the NADH generated in glycolysis. The predictions in case 1 show highly decreased uptake of glucose, yet a small amount of glucose is still taken up, resulting in acetate and lactate production. As discussed in 3.2.1, it is possible that glucose is being taken up *in vivo*, even though this is not detected by measurements, which is in line with model predictions and would also explain the lower growth rate observed *in silico* in case 2 compared to *in vivo*. The higher growth rate *in vivo* compared to *in silico* in case 1 is due to a much higher arginine uptake than measured in the WT. Also, in line with published studies (Chen et al., 2016), addition of glycerol to the *adhE* mutant increases the growth rate to almost WT levels (Figure 4d). Similarly to the WT predictions, the model in case 1 predicts slightly higher growth rate and uptake rates of glucose and glycerol, resulting in higher secretion of by-products. But as before, the flux distribution is very similar to the one measured *in vivo*.

In all four conditions in figure 4 the *in silico* predictions in case 2 and the *in vivo* data are almost identical, with the exception of the few instances described above. In few cases discrepancies can be explained by carbon imbalance *in vivo*, which is most likely due to measurement uncertainties. Taken together, these results show that the model can be used to accurately predict metabolic behavior, without requiring extensive experimental data.

#### 3.2.3 Predicted effects of an active 1,2-propanediol pathway

*L. reuteri* JCM 1112 appears to lack only one enzyme, methylglyoxal synthase (MGS) in the 1,2-propanediol- and 1-propanol biosynthetic pathways (see section 3.1.1). Here we used the model to predict how *mgs* gene insertion would affect the metabolism, specifically in the *adhE* mutant grown on glucose. The mutant grows poorly on glucose due to redox imbalance (section 3.2.2). The synthesis of both 1,2-propanediol and 1-propanol consume NADH and activating these pathways therefore has the potential to restore growth. As in case 1 above, the model was constrained only with experimental uptake rates of glucose and the 5 amino acids from the WT grown on glucose. The *adhE* gene was knocked out *in silico*, and we then compared flux predictions with an added *mgs* (Figure 5) and without it (Figure 4c). The *mgs* addition resulted in a highly increased growth rate (0.11 to 0.49 h^−1^) as well as growth-coupled production of 1-propanol (14.7 mmol gDW^−1^ h^−1^). Given the good agreement between *in silico* predictions and *in vivo* measurements in section 3.2.2, the expression of this gene at a sufficiently high level *in vivo* is expected to result in a relatively fast growing 1-propanol producing cell factory.

**Figure 5.**
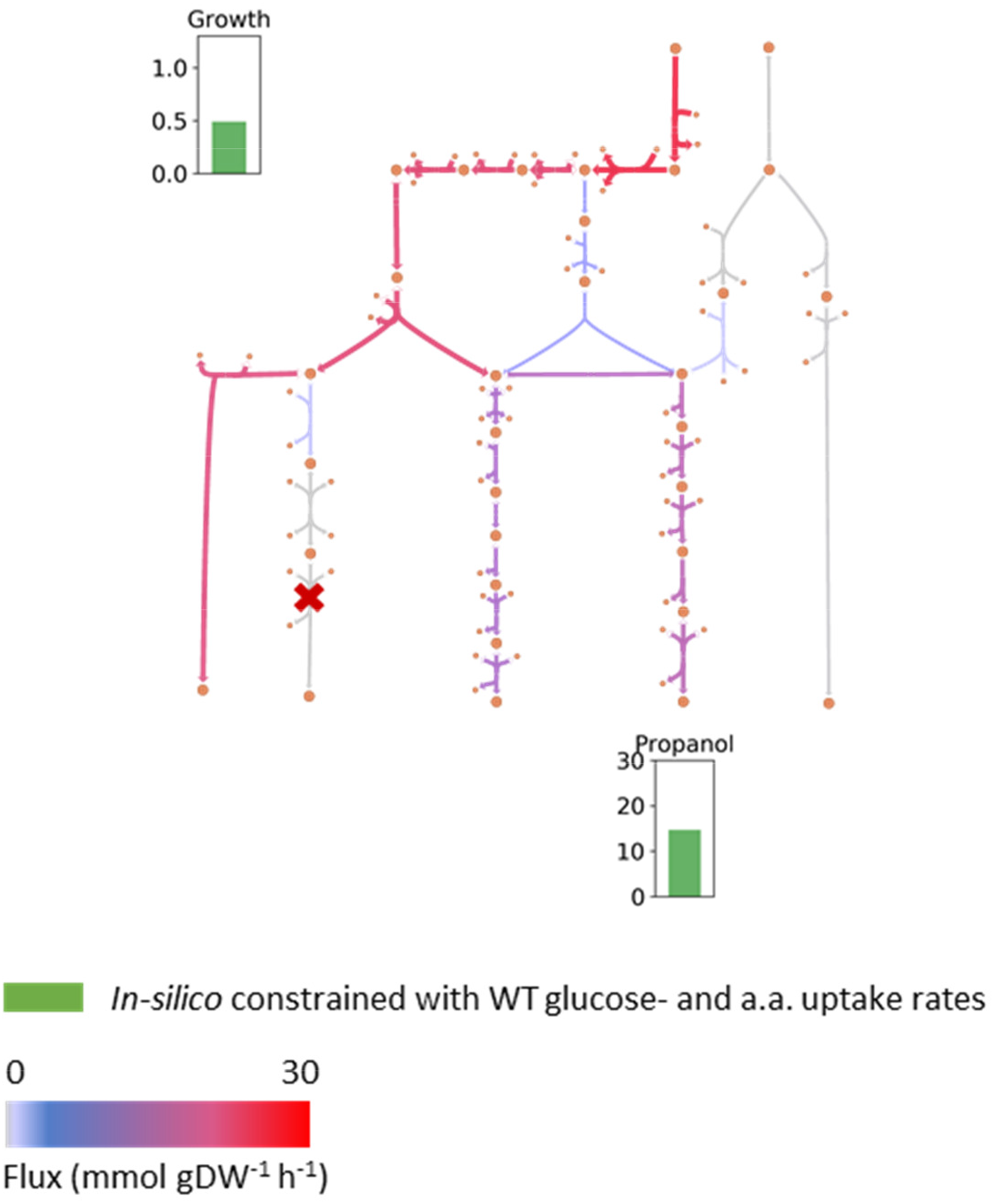
Predicted flux distribution, growth rate and 1-propanol production of adhE mutant grown on glucose, with active 1,2-propanediol and 1-propanol pathways. The model was constrained with average experimental uptake rates of the WT grown on glucose and optimized for growth. Units for propanol secretion rate is in mmol gDW^−1^ h^−1^ and growth rate in h^−1^.

#### 3.2.4 Model-based analysis of *L. reuteri* as a cell factory

LAB are natural producers of several chemicals of industrial interest (Bosma et al., 2017; Papagianni, 2012; Sauer, Russmayer, Grabherr, Peterbauer, & Marx, 2017). They possess high sugar uptake rates and, in many species, the central metabolism is only weakly coupled to biomass formation because of their adaptation to nutrient rich environments. As a result, the carbon source is mostly used for energy gain and is converted to fermentation products in high yields. Combined with high tolerance to environmental stress, these properties have led to significant interest in using LAB as cell factories.

The heterofermentative nature of *L. reuteri* and the dominance of the phosphoketolase over the Embden-Meyerhof-Parnas pathway make some target compounds less suitable than others, with lactic acid being an obvious example. On the other hand, these properties can also be used to an advantage as is demonstrated here We used our newly established *L. reuteri* metabolic model to study the feasibility of this organism to produce some of the compounds that have been the subject of recently published LAB metabolic engineering experiments. These native and non-native compounds include a flavoring compound (acetoin), a food additive (L-alanine), biofuels (1-propanol and ethanol), chemical building blocks (acetaldehyde and 2,3-butanediol) and an environmentally friendly solvent (ethyl lactate). The last compound has recently been produced in an engineered *E. coli* strain (Lee & Trinh, 2018) and is an interesting target in *L. reuteri* since it is a condensation product of the two major products of glucose fermentation via the phosphoketolase pathway, lactate and ethanol.

The suitability of *L. reuteri* for producing a particular compound was assessed in terms of the maximum theoretical yield, using a fixed glucose uptake rate (Table 4). This gives an overly optimistic estimate of product yields in most cases since it completely ignores variations in enzyme efficiency, compound toxicity, regulation and other issues outside the scope of the model. The maximum flux is still useful to identify products that appear to be ill suited for a particular metabolism as well as products that may be suitable.

**Table 4.**
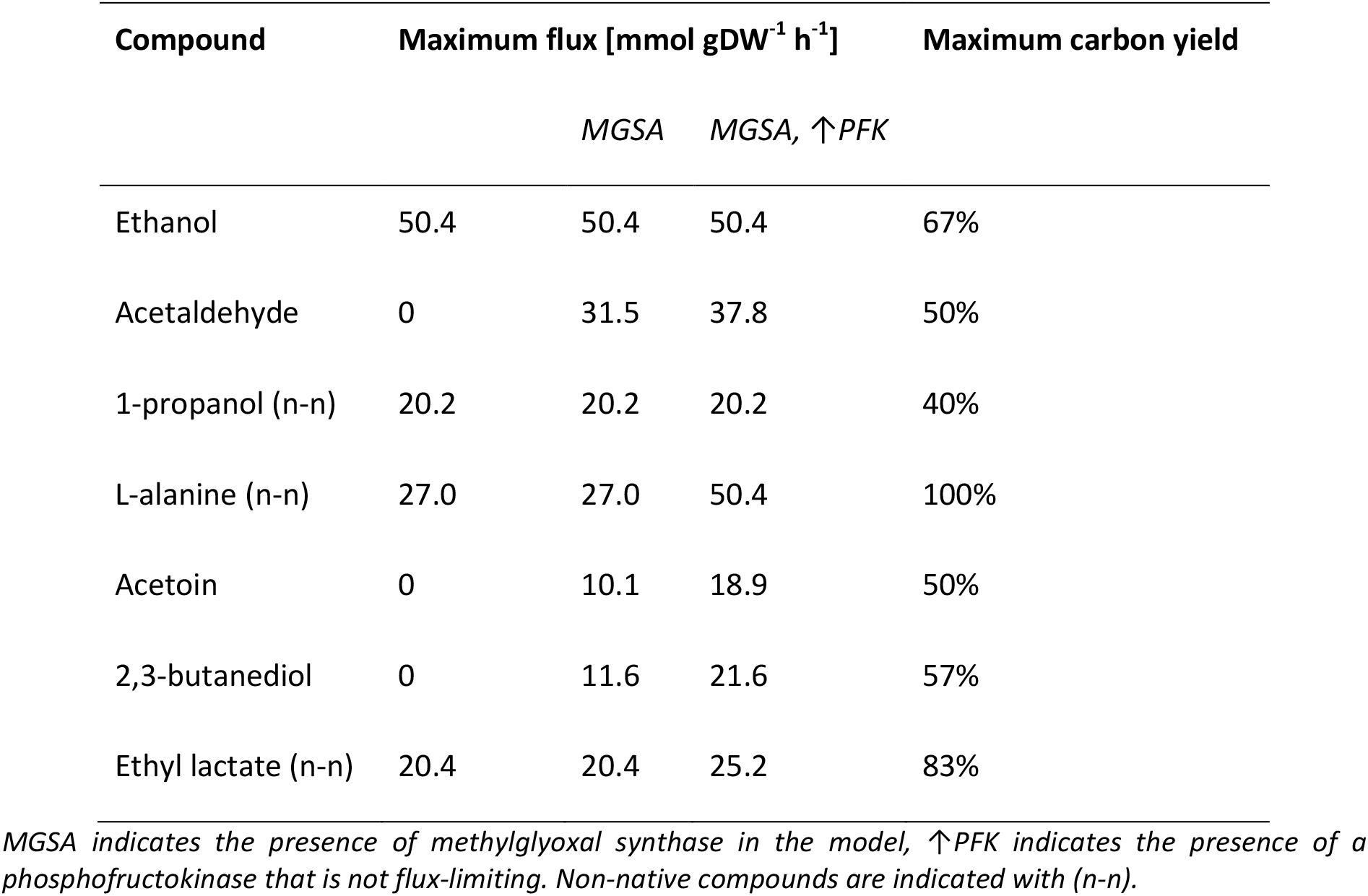
Model predictions of the maximum flux of selected target compounds in L. reuteri assuming a maximum glucose uptake rate of 25.2 mmol gDW^−1^ h^−1^.

The predicted flux for acetaldehyde, acetoin and 2,3-butanediol, which are all derived from acetyl-CoA, was low, suggesting that the metabolism in the wild type is not well suited for overproducing these compounds. The flux increased significantly upon addition of methylglyoxal synthase, suggesting the importance of the 1-propanol pathway in cofactor balancing (section 3.2.3). Addition of glycerol to the medium served the same purpose and increased the predicted flux in all cases (data not shown), which is in line with glycerol being known and used as an external electron sink in *L. reuteri* (Dishisha et al., 2014). For all the compounds except ethanol and 1-propanol, the addition of a fully functional phosphofructokinase was predicted to increase the yields even further (Table 4). Such a strategy has been shown successful for mannitol production (Papagianni & Legiša, 2014).

Taken together, the model suggests that *L. reuteri* is better suited for producing compounds derived from pyruvate than compounds derived from acetyl-CoA and that the simultaneous expression of heterologous MGSA and PFK enzymes is a general metabolic engineering strategy for increasing product yields in *L. reuteri*.

## 4. Conclusions

In this study, we have established a manually curated genome-scale metabolic model of *L. reuteri* JCM 1112, referred to as Lreuteri_530, and validated it with experimental data. We identified several knowledge gaps in the metabolism of this organism that we resolved with a combination of experimentation and modeling. The distribution of flux between the PKP and EMPP pathways is strain-specific and in line with other studies, we found that the EMPP activity is maximally around 7% of total glycolytic flux during early exponential phase. The predictive accuracy of the model was estimated by comparing predictions with experimental data. Several scenarios were tested both *in vivo* and *in silico*, including addition of glycerol to a glucose-based growth medium and the deletion of the *adhE* gene, which encodes a bifunctional aldehyde/alcohol dehydrogenase. The results showed that the model gives accurate predictions, both with respect to growth rate and uptake- and secretion rates of main metabolites in the central metabolism. This indicates that the model can be useful for predicting metabolic engineering strategies, such as growth-coupled production of 1-propanol. The model also serves as a starting point for the modeling of other L. *reuteri* strains and related species. The model is available in SBML, Matlab and JSON formats at https://github.com/steinng/reuteri as well as in Additional file 6. Metabolic maps in Escher format are provided in Additional file 4. The Escher maps together with the model in JSON format can be used directly with the Escher-FBA online tool (Rowe, Palsson, & King, 2018) as well as the Caffeine cell factory design and analysis platform (https://caffeine.dd-decaf.eu/).

## 5. Declarations

### Ethics approval and consent to participate

Not applicable.

### Consent for publication

Not applicable.

### Availability of data and material

The model, experimental data, code and other relevant material are available from github.com/steinng/reuteri and Additional files.

## Authors’ contributions

TK and SG curated and validated the metabolic reconstruction, performed numerical simulations and wrote the manuscript. EFB performed all experimental work except the bioreactor cultivations, performed data processing and analysis, curated the metabolic reconstruction and wrote the manuscript. FBdS constructed the draft model, performed bioreactor cultivations, analyzed the resulting data and revised the manuscript. EÖ curated the original draft metabolic reconstruction, processed and analyzed the genome sequencing data and revised the manuscript. ATN and MJH were involved in the metabolic reconstruction and revised the manuscript. BSF and LF performed a bioreactor cultivation, analyzed the resulting data and revised the manuscript. EFB, TK, SG and ATN conceived and coordinated this study. All authors read and approved the final manuscript.

## Acknowledgements

The authors would like to thank Bjarke Krysel Christensen, Steen Troels Jørgensen and Brian Kobmann from Novozymes for providing strain SJ11774; Anna Koza from DTU Biosustain for performing the genome sequencing; Amalie Melton Axelsen from DTU Biosustain for technical support with construction and analysis of the *adhE* mutant strain.

