## Supplementary material for "A metabolic reconstruction of *Lactobacillus reuteri* JCM 1112 and analysis of its potential as a cell factory": Additional_file_3a.ipynb

```
# %% [markdown]
# This notebook:
### 1. Reads the SBML model and adds tailored constraint
### 2. Reads in neccessary experimental data file
### 3. Creates bar plot in section 3.2.1
### 4. Creates small bar plots and flux files (for Escher), used to create figures in section 3.2.2 and 3.2.3
### 5. Predicts theoretical maximum yields of target compounds, with and without added engineering strategies in section 3.2.4

# %% [markdown]
### Import

# %% [code]
import cobra
from cobra import Reaction
import numpy as np
import matplotlib.pyplot as plt

# %% [markdown]
### Add local paths to neccessary files

# %% [code]
sbml_model = '' # Add path to model here
exp_data_file = '' # Add path to experimental data file here
path_to_output_flux_files = '' # Add path to folder where you want flux files to be saved to

# %% [markdown]
### Functions used in this notebook

# %% [code]
def load_sbml_reuteri_model(model):
    model = cobra.io.read_sbml_model(model)
    EMP_PKP_flux = model.problem.Constraint(model.reactions.PFK.flux_expression*0.93 - model.reactions.G6PDH2r.flux_expression*0.07, lb=-1000, ub=0)
    model.add_cons_vars(EMP_PKP_flux)
    return model

def constrain_and_optimize(data_dictionary):
    for reaction, rate in data_dictionary.items():
        if reaction == 'BIOMASS':
            Experimental_list.append(float(rate))
            continue
        if float(rate) <= 0:
            model.reactions.get_by_id(reaction).lower_bound = float(rate)
            model.reactions.get_by_id(reaction).upper_bound = 0
        if float(rate) >= 0:
            model.reactions.get_by_id(reaction).upper_bound = float(rate)
            model.reactions.get_by_id(reaction).lower_bound = 0
        else:
            continue
    sol = model.optimize()
    Predicted_list.append(sol.f)
    return Predicted_list, Experimental_list

def create_flux_file(optimization_solution, name_of_output_flux_file=''):
    Flux_txt = open(path_to_output_flux_files+'/'+name_of_output_flux_file+'.txt','w')
    Flux_txt.write('rxn, flux\n')
    for y in range(len(model.reactions)):
        if model.reactions[y].id == 'QUIND':
            Flux_txt.write(model.reactions[y].id + ', ' + str(30) + '\n')
        else:
            Flux_txt.write(model.reactions[y].id + ', ' + str(optimization_solution.x[y]) + '\n')
    Flux_txt.close()
    return

def plot_for_escher(Data_Dictionary, Sol_Vector, Real_Sol_Vector, Reaction_ID, Metabolite_Name, Save_Name='', Strain_Cond=''):
    n_groups = 1
    plt.figure(figsize=(1.5,2))
    number = np.arange(n_groups)
    bar_width = 0.35
    plt.bar(number, abs(Sol_Vector.x[model.reactions.index(Reaction_ID)]), bar_width, alpha=0.6, color='g')
    plt.bar(number + bar_width+0.04, abs(Real_Sol_Vector.x[model.reactions.index(Reaction_ID)]), bar_width, alpha=0.2, color='g')
    plt.bar(number + 2*bar_width+0.08, abs(Data_Dictionary[Reaction_ID]), bar_width, alpha=0.2, color='tab:orange')
    plt.title(Metabolite_Name, fontsize=16)
    plt.ylim(0,30)
    plt.yticks(fontsize=16)
    plt.xticks([])
    plt.tight_layout()
    print(Strain_Cond+Save_Name)
    plt.show()
    plt.close()
    return

def max_flux_of_target(model, EX_objective='', target=''):
    with model:
        model.objective = model.reactions.get_by_id(EX_objective)
        model.add_cons_vars(EMP_PKP_flux)
        sol = model.optimize().f
        model.add_reactions([Reaction('MGS')]) # Add heterologous Methylglyoxal Synthase reaction
        model.reactions.MGS.reaction = 'dhap_c -> mthgxl_c + pi_c'
        MGS_sol = model.optimize().f
    with model:
        model.objective = model.reactions.get_by_id(EX_objective)
        model.reactions.PFK.upper_bound = 1000 # Allow free PFK flux
        model.add_reactions([Reaction('MGS')]) # Add heterologous Methylglyoxal Synthase reaction
        model.reactions.MGS.reaction = 'dhap_c -> mthgxl_c + pi_c'
        MGS_PFK_sol = model.optimize().f
    print(target, '\t', round(sol, 1), '\t', round(MGS_sol, 1), '\t', round(MGS_PFK_sol, 1))

# %% [markdown]
### Read in model and add tailored constraint, which limits flux through EMP pathway to max 7%

# %% [code]
model = load_sbml_reuteri_model(sbml_model)

# %% [markdown]
### Read in experimental data file and create data dictionaries

# %% [code]
WT_glc_dict = {}
WT_glc_glyc_dict = {}
SJ_glc_dict = {}
SJ_glc_glyc_dict = {}
adhE_glc_dict = {}
adhE_glc_glyc_dict = {}

data_file = open(exp_data_file, 'r')

for line in data_file.readlines()[1:]:
    line = line.split('\t')
    WT_glc_dict[line[0]] = float(line[2])
    WT_glc_glyc_dict[line[0]] = float(line[3])
    SJ_glc_dict[line[0]] = float(line[4])
    SJ_glc_glyc_dict[line[0]] = float(line[5])
    adhE_glc_dict[line[0]] = float(line[6])
    adhE_glc_glyc_dict[line[0]] = float(line[7].strip('\n'))
data_file.close()

# %% [markdown]
### Constrain model, optimize and create lists for bar plot in section 3.2.1 and variables for section 3.2.2

# %% [code]
Predicted_list = []
Experimental_list = []

constrain_and_optimize(WT_glc_dict) # For section 3.2.1
constrain_and_optimize(WT_glc_glyc_dict) # For section 3.2.1
constrain_and_optimize(SJ_glc_dict) # For section 3.2.1
SJ_glc_real_sol = cobra.flux_analysis.pfba(model) # For section 3.2.2
constrain_and_optimize(SJ_glc_glyc_dict) # For section 3.2.1
SJ_glc_glyc_real_sol = cobra.flux_analysis.pfba(model) # For section 3.2.2
constrain_and_optimize(adhE_glc_dict) # For section 3.2.1
with model:
    model.reactions.ACALD.knock_out()
    adhE_glc_real_sol = cobra.flux_analysis.pfba(model) # For section 3.2.2
constrain_and_optimize(adhE_glc_glyc_dict) # For section 3.2.1
with model:
    model.reactions.ACALD.knock_out()
    adhE_glc_glyc_real_sol = cobra.flux_analysis.pfba(model) # For section 3.2.2

# %% [markdown]
### Create bar plot in section 3.2.1

# %% [code]
Name_string = ('A', 'B', 'C', 'D', 'E', 'F')

n_groups = 6

fig, ax = plt.subplots(figsize=(6,3))

index = np.arange(n_groups)
bar_width = 0.35

# Experimental growth rates for all replicates
Replicates = [0,0,0,1,1,2,2,2,3,3,3,4,4,4,5,5,5]
Experimental_Growth_Rates = [0.57,0.68,0.62,0.74,0.69,0.69,0.72,0.68,0.66,0.60,0.57,0.30,0.37,0.38,0.58,0.53,0.52]

# Plot average experimental growth rate - light blue bars
ax.bar(index, Experimental_list, bar_width, alpha=0.4, color='b', label='Experimental growth rate')

# Plot experimental growth rates for each replicate used to determine average - dark blue dots
ax.plot(Replicates, Experimental_Growth_Rates, 'bo', markersize=3)

# Plot predicted growth rates, when model constrained with average experimental values
ax.bar(index + bar_width, Predicted_list, bar_width, alpha=0.4, color='r', label='Predicted growth rate')

ax.set_ylabel('Growth rate ($h^{-1}$)', fontsize=12) 

ax.set_xticks(index + bar_width / 2)
ax.set_xticklabels(Name_string, fontsize=12)
ax.set_ylim(0,1.1)
ax.legend()

fig.tight_layout()
plt.show()

# %% [markdown]
### Create flux files for Escher used in section 3.2.2

# %% [code]
# Model constrained with uptake rates of glucose and 5 amino acids of the type strain (SJ) grown on glucose
for reaction, rate in SJ_glc_dict.items():
    if reaction == 'BIOMASS':
        continue
    if float(rate) <= 0:
        model.reactions.get_by_id(reaction).lower_bound = float(rate)
        model.reactions.get_by_id(reaction).upper_bound = 0
    if float(rate) >= 0:
        model.reactions.get_by_id(reaction).upper_bound = 1000
        model.reactions.get_by_id(reaction).lower_bound = 0
    else:
        continue

model.reactions.EX_lac__D_e.upper_bound = 0
        
# Optimize for growth and create flux file - Wild type (SJ) on glucose
SJ_glc_sol = cobra.flux_analysis.pfba(model) 
create_flux_file(SJ_glc_sol, name_of_output_flux_file='Flux_file_SJ_glc')

# Optimize for growth and create flux file - adhE mutant on glucose
with model:
    model.reactions.ACALD.knock_out()
    AdhE_glc_sol = cobra.flux_analysis.pfba(model)
    create_flux_file(AdhE_glc_sol, name_of_output_flux_file='Flux_file_adhE_glc')

# Optimize for growth and create flux file - Wild type (SJ) on glucose and glycerol
model.reactions.EX_glyc_e.lower_bound = -25
SJ_glc_glyc_sol = cobra.flux_analysis.pfba(model)
create_flux_file(SJ_glc_glyc_sol, name_of_output_flux_file='Flux_file_SJ_glc_glyc')

# Optimize for growth and create flux file - adhE mutant on glucose and glycerol
with model:
    model.reactions.ACALD.knock_out()
    AdhE_glc_glyc_sol = cobra.flux_analysis.pfba(model)
    create_flux_file(AdhE_glc_glyc_sol, name_of_output_flux_file='Flux_file_adhE_glc_glyc')

# %% [markdown]
### Create small bar plots for section 3.2.2

# %% [code]
Reaction_IDs = ['EX_glc__D_e', 'EX_glyc_e', 'EX_lac__L_e', 'EX_ac_e', 'EX_etoh_e', 'EX_13ppd_e', 'BIOMASS']
Metabolites_Names = ['Glucose', 'Glycerol', 'Lactate', 'Acetate', 'Ethanol', '1,3-ppd', 'Growth']
Save_Names = ['__glc', '__glyc', '__lac', '__ac', '__etoh', '__13ppd', '__GR']

for index in range(len(Reaction_IDs)):
    plot_for_escher(SJ_glc_dict, SJ_glc_sol, SJ_glc_real_sol, Reaction_IDs[index], Metabolites_Names[index], Save_Name=Save_Names[index], Strain_Cond='SJ_glc')

for index in range(len(Reaction_IDs)):
    plot_for_escher(SJ_glc_glyc_dict, SJ_glc_glyc_sol, SJ_glc_glyc_real_sol, Reaction_IDs[index], Metabolites_Names[index], Save_Name=Save_Names[index], Strain_Cond='SJ_glc_glyc')
    
for index in range(len(Reaction_IDs)):
    plot_for_escher(adhE_glc_dict, AdhE_glc_sol, adhE_glc_real_sol, Reaction_IDs[index], Metabolites_Names[index], Save_Name=Save_Names[index], Strain_Cond='AdhE_glc')
    
for index in range(len(Reaction_IDs)):
    plot_for_escher(adhE_glc_glyc_dict, AdhE_glc_glyc_sol, adhE_glc_glyc_real_sol, Reaction_IDs[index], Metabolites_Names[index], Save_Name=Save_Names[index], Strain_Cond='AdhE_glc_glyc')

# %% [markdown]
### Create flux file and bar plots for section 3.2.3

# %% [code]
model.reactions.EX_glyc_e.lower_bound = 0 # Reset uptake rate of glycerol to 0

with model:
    model.reactions.ACALD.knock_out()
    MGS = Reaction('MGS')
    model.add_reactions([MGS])
    model.reactions.MGS.reaction = 'dhap_c  -> mthgxl_c + pi_c'
    GR_ppoh = model.optimize().f
    sol_ppoh = model.optimize()
    plt.figure(figsize=(1.3,2))
    plt.bar(2, sol_ppoh.x[model.reactions.index('EX_ppoh_e')], color='g', alpha=0.6, width=1.1)
    plt.title('Propanol', fontsize=16)
    plt.xlim(1,3)
    plt.ylim(0,30)
    plt.yticks(fontsize=16)
    plt.xticks([])
    plt.tight_layout()
    print('Propanol produced when optimized for growth with an active 1,2-propanediol pathway:')
    plt.show()
    plt.close()
    create_flux_file(sol_ppoh, name_of_output_flux_file='Flux_file_adhE-glc_with_ppoh_pathway')
    plt.figure(figsize=(1.4,2))
    plt.bar(2, GR_ppoh, color='g', alpha=0.6, width=1.1)
    plt.title('Growth', fontsize=16)
    plt.ylim(0,1.3)
    plt.xlim(1,3)
    plt.yticks(fontsize=16)
    plt.xticks([])
    plt.tight_layout()
    print('Growth rate with an active 1,2-propanediol pathway:')
    plt.show()
    plt.close()


# %% [markdown]
### Predict production rate of target metabolites, with and without engineering strategies for section 3.2.4

# %% [code]
# This is the default model (with AA and nucleotide uptake) but without limited PFK flux constraint)
model = cobra.io.read_sbml_model(sbml_model)

# PFK limiting flux constraint
EMP_PKP_flux = model.problem.Constraint(model.reactions.PFK.flux_expression*0.93 - model.reactions.G6PDH2r.flux_expression*0.07, lb=-1000, ub=0)

# Allow free secretion of by-products
model.reactions.EX_ac_e.upper_bound = 1000
model.reactions.EX_btd_RR_e.upper_bound = 1000
model.reactions.EX_13ppd_e.upper_bound = 1000
model.reactions.EX_lac__L_e.upper_bound = 1000
model.reactions.EX_lac__D_e.upper_bound = 1000
model.reactions.EX_etoh_e.upper_bound = 1000

# Don´t allow uptake of amino acids and nucleotides. We only want to predict production of products from glucose.
model.reactions.EX_glu__L_e.lower_bound = 0
model.reactions.EX_arg__L_e.lower_bound = 0
model.reactions.EX_ser__L_e.lower_bound = 0
model.reactions.EX_asn__L_e.lower_bound = 0
model.reactions.EX_asp__L_e.lower_bound = 0
model.reactions.EX_ade_e.lower_bound = 0
model.reactions.EX_ura_e.lower_bound = 0
model.reactions.EX_xan_e.lower_bound = 0
model.reactions.EX_gua_e.lower_bound = 0

# Ethonal
max_flux_of_target(model, EX_objective='EX_etoh_e', target='Ethanol')

# Acetaldehyde
max_flux_of_target(model, EX_objective='EX_acald_e', target='Acetaldehyde')

# 1-propanol (n-n)
with model:
    model.add_reactions([Reaction('MGS')]) # Add heterologous Methylglyoxal Synthase reaction
    model.reactions.MGS.reaction = 'dhap_c -> mthgxl_c + pi_c'
    max_flux_of_target(model, EX_objective='EX_ppoh_e', target='1-Propanol (n-n)')

# Alanine (n-n)    
with model:
    model.add_reactions([Reaction('ALAD_L')]) # Add heterologous Alanine dehydrogenase reaction
    model.reactions.ALAD_L.reaction = 'ala__L_c + h2o_c + nad_c <=> h_c + nadh_c + nh4_c + pyr_c'
    max_flux_of_target(model, EX_objective='EX_ala__L_e', target='L-Alanine (n-n)')

# Acetoin
max_flux_of_target(model, EX_objective='EX_actn__R_e', target='R-Acetoin')

# 2,3-butanediol
max_flux_of_target(model, EX_objective='EX_btd_RR_e', target='2,3-butanediol')

# Ethyl lactate (n-n)
with model:
    model.add_reactions([Reaction('LACAT')]) # Add heterologous Lacatate Acyltransferase reaction
    model.reactions.LACAT.reaction = 'ac_c + lcoa__L_c <=> accoa_c + lac__L_c'
    model.add_reactions([Reaction('VAAT')]) # Add heterologous Esterification reaction
    model.reactions.VAAT.reaction = 'etoh_c + lcoa__L_c <=> ethyllac_c + coa_c'
    model.add_reactions([Reaction('DM_ethyllac_c')]) # Add Demand reaction for Ethyl lactate
    model.reactions.DM_ethyllac_c.reaction = 'ethyllac_c -> '
    max_flux_of_target(model, EX_objective='DM_ethyllac_c', target='Ethyl lactate (n-n)')


# %% [code]
```
