## Supplementary material for "A metabolic reconstruction of *Lactobacillus reuteri* JCM 1112 and analysis of its potential as a cell factory": Additional_file_7.docx

**Biomass composition of *L. reuteri***

Protein:DNA:RNA:Lipid:LTA:CPS:PG ratio of biomass is not known for reuteri. Lipoteichoic acid (LTA) from *L. lactis* (Oliveira, Nielsen, & Förster, 2005) is used because *L. plantarum* also produces Wall teichoic acid (WTA), which *L. reuteri* and *L. lactis* do not. Peptidoglycan in *L. plantarum* (Teusink et al., 2006) was calculated from *Bacillus cereus*, so here peptidoglycan from *L. lactis* is used. Protein ratio was measure for *L. reuteri* in this study. Otherwise numbers from *L. plantarum* were used for the macromolecular composition.

| **Component** | | **Fraction (% w/w)** | **Norm Fraction (% w/w)** | **Molar mass (g mol^-1^)** | **Coefficent (mmol gDW^-1^)** | **Source** |
| --- | --- | --- | --- | --- | --- | --- |
| Protein | PROT_LRE_c | 35,4% | 43,0% | 129,9 | 3,311 | *L. reuteri* (This study) |
| DNA | DNA_LRE_c | 1,9% | 2,3% | 309,5 | 0,075 | *L. plantarum* (Oliveira et al., 2005; Teusink et al., 2006) |
| RNA | RNA_LRE_c | 9,0% | 10,9% | 321,6 | 0,340 | *L. plantarum* (Teusink et al., 2006) |
| Lipids | LIP_LRE_c | 6,3% | 7,7% | 795,1 | 0,096 | *L. plantarum* (Teusink et al., 2006) |
| Lipoteichoic acid | LTAtotal_LRE_c | 8,0% | 9,7% | 5290,4 | 0,018 | *L. lactis* (Oliveira et al., 2005) |
| Polysaccharides | CPS_LRE_c | 9,9% | 12,0% | 648 | 0,186 | *L. plantarum* (Teusink et al., 2006) |
| Peptidoglycan | PGlac2_c | 11,8% | 14,3% | 992 | 0,145 | *L. lactis* (Oliveira et al., 2005) |
|  | Total | 82,3% | 100% |  |  |  |

0.186 CPS_LRE_c + 0.075 DNA_LRE_c + 0.096 LIP_LRE_c + 0.018 LTAtotal_LRE_c + 0.145 PGlac2_c + 3.311 PROT_LRE_c + 0.340 RNA_LRE_c + 10.2 atp_c + 1e-05 btn_c + 0.0002 coa_c + 10.2 h2o_c + 0.0002 nad_c + 1e-06 pydx5p_c + 1e-05 thf_c + 1e-05 thmpp_c + 0.0002 udcpdp_c + 0.00001 adeadocbl_c --> 10.2 adp_c + 10.2 h_c + 10.2 pi_c

Flux through ATPM is fixed at:  0.36 mmol h^-1^ gDW^-1^ (*L. plantarum* (Teusink et al., 2006))

**Protein**

Amino acid composition analyzed following cell pellet hydrolysis (Ansynth).

| **Amino Acid** | **Fraction g/g_total a.a._** | **Fraction g/gDW** | **Fraction g/gDW (only protein)** | **MW (g/mol)** | **Fraction mole/gDW** | **Molar ratio** |
| --- | --- | --- | --- | --- | --- | --- |
| Ala | 10,4% | 5,2% | 0,2% | 89,1 | 2,25E-05 | 0,7%* |
| Arg | 4,7% | 2,4% | 2,4% | 174,2 | 1,35E-04 | 4,1% |
| Asp + Asn | 13,4% | 6,7% | 6,7% | 132,1 | 5,08E-04 |  |
| Asp |  |  |  |  |  | 7,7% |
| Asn |  |  |  |  |  | 7,7% |
| Cys | 0,8% | 0,4% | 0,4% | 121,2 | 3,15E-05 | 1,0% |
| Glu + Gln | 13,8% | 6,9% | 4,8% | 146,6 | 3,27E-04 |  |
| Glu |  |  |  |  |  | 2,7%* |
| Gln |  |  |  |  |  | 7,1% |
| Gly | 5,2% | 2,6% | 2,6% | 75,1 | 3,50E-04 | 10,6% |
| His | 2,1% | 1,1% | 1,1% | 155,2 | 6,93E-05 | 2,1% |
| Ile | 4,9% | 2,5% | 2,5% | 131,1 | 1,89E-04 | 5,7% |
| Leu | 7,3% | 3,7% | 3,7% | 131,1 | 2,81E-04 | 8,5% |
| Lys | 9,4% | 4,7% | 4,7% | 146,2 | 3,23E-04 | 9,8% |
| Met | 2,7% | 1,3% | 1,3% | 149,2 | 8,93E-05 | 2,7% |
| Phe | 3,5% | 1,8% | 1,8% | 165,2 | 1,08E-04 | 3,3% |
| Pro | 3,0% | 1,5% | 1,5% | 115,1 | 1,33E-04 | 4,0% |
| Ser | 3,8% | 1,9% | 1,9% | 105,1 | 1,80E-04 | 5,4% |
| Thr | 5,1% | 2,5% | 2,5% | 119,1 | 2,13E-04 | 6,4% |
| Trp | 0,9% | 0,5% | 0,5% | 204,2 | 2,31E-05 | 0,7% |
| Tyr | 3,4% | 1,7% | 1,7% | 181,2 | 9,30E-05 | 2,8% |
| Val | 5,4% | 2,7% | 2,7% | 117,2 | 2,33E-04 | 7,0% |
| **Total** | **100%** | **50,2%** | **43,0%** | **129,9** | **3,31E-03** | **100%** |

*Glutamate and alanine in lipoteichoic acid and peptidoglycan are subtracted from measured values.

Alanine in lipoteichoic acid:

| mmol LTA / gDW | 0,018 |
| --- | --- |
| mol alanine / mol LTA | 15,3 |
| mol alanine in LTA / gDW | 0,0003 |
| Alanine molar mass (g/mol) | 89,094 |
| **g alanine in LTA / gDW** | **0,0245** |

Alanine in peptidoglycan:

| mmol PGlac2 / gDW | 0,145 |
| --- | --- |
| mol alanine / mol LTA | 2 |
| mol alanine in LTA / gDW | 0,0003 |
| Alanine molar mass (g/mol) | 89,094 |
| **g alanine in LTA / gDW** | **0,0258** |

Glutamate in Peptidoglycan:

| mmol PGlac2 / gDW | 0,145 |
| --- | --- |
| mol glutamate / mol LTA | 1 |
| mol glutamate in LTA / gDW | 0,0002 |
| Glutamate molar mass (g/mol) | 147,13 |
| **g glutamate in LTA / gDW** | **0,0213** |

**Protein reaction:**
0.306 atp_c + 2 gtp_c + 2.306 h2o_c + 0.0062 alatrna_c + 0.0412 argtrna_c + 0.0775 asntrna_c + 0.0775 asptrna_c + 0.0096 cystrna_c + 0.0721 glntrna_c + 0.0189 glutrna_c + 0.1068 glytrna_c + 0.0211 histrna_c + 0.0578 iletrna_c + 0.0857 leutrna_c + 0.0986 lystrna_c + 0.0272 mettrna_c + 0.0329 phetrna_c + 0.0404 protrna_c + 0.0548 sertrna_c + 0.0650 thrtrna_c + 0.0070 trptrna_c + 0.0284 tyrtrna_c + 0.0711 valtrna_c --> PROT_LRE_c + 0.306 adp_c + 2.0 gdp_c + 2.306 h_c + 2.306 pi_c + 0.0062 trnaala_c + 0.0412 trnaarg_c + 0.0775 trnaasn_c + 0.0775 trnaasp_c + 0.0096 trnacys_c + 0.0910 trnaglu_c + 0.1068 trnagly_c + 0.0211 trnahis_c + 0.0578 trnaile_c + 0.0857 trnaleu_c + 0.0986 trnalys_c + 0.0272 trnamet_c + 0.0329 trnaphe_c + 0.0404 trnapro_c + 0.0548 trnaser_c + 0.0650 trnathr_c + 0.0070 trnatrp_c + 0.0284 trnatyr_c + 0.0711 trnaval_c

**DNA**

Calculated from JCM1112 genome

| **Nucleotide** | **Ratio** | **MW (g mol^-1^)** | **MW * Ratio (g mol^-1^)** |
| --- | --- | --- | --- |
| dAMP | 0,30560617 | 331,2 | 101,2 |
| dTMP | 0,30560617 | 304,2 | 93,0 |
| dCMP | 0,19439383 | 289,2 | 56,2 |
| dGMP | 0,19439383 | 304,2 | 59,1 |
| **Average MW of DNA** | | | **309,5** |

**DNA reaction:**
1.37 atp_c + 0.310284595735 datp_c + 0.180906354753 dctp_c + 0.207484087701 dgtp_c + 0.301324961811 dttp_c + 1.37 h2o_c --> DNA_LRE_c + 1.37 adp_c + 1.37 h_c + 1.37 pi_c + ppi_c

**RNA**

Calculated from JCM1112 genome, assuming equal transcription

| **Nucleotide** | **Ratio** | **MW (g mol^-1^)** | **MW * Ratio (g mol^-1^)** |
| --- | --- | --- | --- |
| AMP | 0,31434679 | 329,2 | 103,5 |
| UMP | 0,29106444 | 306,2 | 89,1 |
| CMP | 0,18074957 | 305,2 | 55,2 |
| GMP | 0,21383921 | 345,2 | 73,8 |
| **Average MW of RNA** | | | **321,6** |

**RNA reaction:**
0.710810743235 atp_c + 0.187747387186 ctp_c + 0.21245220899 gtp_c + 0.288989660589 utp_c --> RNA_LRE_c + 0.4 adp_c + 0.4 h_c + 0.4 pi_c + ppi_c

**Lipids** (Liu, Hou, Zhang, Zeng, & Qiao, 2014)

| **Fatty acid** | **Ratio** | **MW (g mol^-1^)** | **Formula** | **MW * Ratio (g mol^-1^)** |
| --- | --- | --- | --- | --- |
| C14:0 | 0,0687 | 228,38 | C_14_H_28_O_2_ | 15,68971 |
| C16:0 | 0,2566 | 256,43 | C_16_H_32_O_2_ | 65,79994 |
| C16:1 | 0,0258 | 254,41 | C_16_H_30_O_2_ | 6,563778 |
| C18:0 | 0,075 | 284,48 | C_18_H_36_O_2_ | 21,336 |
| C18:1 | 0,1591 | 282,47 | C_18_H_34_O_2_ | 44,94098 |
| C18:2 | 0,3519 | 280,45 | C_18_H_32_O_2_ | 98,69036 |
| C18:3 | 0,0418 | 278,44 | C_18_H_30_O_2_ | 11,63879 |
| cyc19:0 | 0,0211 | 296,49 | C_19_H_36_O_2_ | 6,255939 |
| **Average MW of fatty acids** | | | | **270,9155** |

**Fatty acids:**
agly3p_LRE_c + 0.0211 cpocdacp_c + 0.0258 hdeACP_c + 0.075 ocdacp_c + 0.0418 ocdctrACP_c + 0.3519 ocdcyaACP_c + 0.1591 octeACP_c + 0.2566 palmACP_c + 0.0687 tdeacp_c --> ACP_c + pa_LRE_c

Phospholipids ratio from *L. plantarum* used here, as no *L. reuteri* data was available

| **Phospholipid** | **Ratio** | **MW with average fatty acid (g mol^-1^)** | **MW * Ratio (g mol^-1^)** |
| --- | --- | --- | --- |
| Phosphatidylglycerol | 0,75 | 751,95 | 563,96 |
| 1-lysyl phosphatidylglycerol | 0,23 | 882,14 | 202,89 |
| Cardiolipin | 0,02 | 1411,81 | 28,24 |
| **Average MW of phospholipids** | | | **795,1** |

**Lipid reaction:**
0.02 clpn_LRE_c + 0.23 lyspg_LRE_c + 0.75 pg_LRE_c --> LIP_LRE_c

**Lipoteichoic acid** (Bron, Baarlen, & Kleerebezem, 2011; Kleerebezem et al., 2010; Walter et al., 2007)

20 glycerol phosphate (Gro-P) residues

Gro-P residues were substituted with d-alanyl esters (74-79%, average 76,5%)

Gro-P residues were substituted with glycosyl residues (6%)

| **Component** | **Average molar ratio** | **Molar mass (g mol^-1^)** |
| --- | --- | --- |
| Diglucosyl diacylglycerol | 1 | 906,3 |
| Glycerol phosphate | 20 | 154,0 |
| D-alanine | 15,3 | 71,1 |
| D-glucose | 1,2 | 180,2 |
| **Average molar mass of Lipoteichoic acid** | | **5290,4** |

**LTA reaction:**
0.175 LTA_LRE_c + 0.765 LTAala_LRE_c + 0.06 LTAglc_LRE_c --> LTAtotal_LRE_c

L. reuteri does not produce Wall teichoic acid

**Polysaccharides** (Ksonzeková et al., 2016)

Glucan homopolysaccharides (merely glucose units)

D-glucose, 4 units, 162 g mol^-1^ per unit 🡪 648 g mol^-1^

**Polysaccharide reaction:**
4.0 h2o_c + 4.0 udpg_c <=> CPS_LRE_c + 5.0 h_c + 3.0 udp_c + ump_c

**Peptidoglycan**

Assumed the same as *L.* *plantarum*

**Peptidoglycan reaction:**
uaagmdalac_c --> PGlac2_c + udcpdp_c

**Glycogen**

Glycogen in *B. subtilis* biomass was negligible (Dauner & Sauer, 2001; Dauner, Storni, & Sauer, 2001). Assumed the same here.

**ATP requirements**

No data for *L. reuteri* available. NGAM assumed the same as *L. plantarum.* GAM calculated by constraining model with experimental uptake- and secretion rates of metabolites, and growth rate of *L. reuteri*.

**Ref**

Bron, P. A., Baarlen, P. Van, & Kleerebezem, M. (2011). Emerging molecular insights into the interaction between probiotics and the host intestinal mucosa. *Nature Reviews Microbiology*, *10*(1), 66–78. https://doi.org/10.1038/nrmicro2690

Dauner, M., & Sauer, U. (2001). Stoichiometric Growth Model for Riboflavin-Producing Bacillus subtilis. *Biotechnology and Bioengineering*, *76*(2), 132–143.

Dauner, M., Storni, T., & Sauer, U. W. E. (2001). Bacillus subtilis Metabolism and Energetics in Carbon-Limited and Excess-Carbon Chemostat Culture. *Journal of Bacteriology*, *183*(24), 7308–7317. https://doi.org/10.1128/JB.183.24.7308

Kleerebezem, M., Hols, P., Bernard, E., Rolain, T., Zhou, M., Roland, J., & Bron, P. A. (2010). The extracellular biology of the lactobacilli. *FEMS Microbiology Reveiws*, *34*, 199–230. https://doi.org/10.1111/j.1574-6976.2009.00208.x

Ksonzeková, P., Bystricky, P., Vlcková, S., Pätoprst, V., Pulzová, L., Mudronová, D., … Tkáciková, L. (2016). Exopolysaccharides of Lactobacillus reuteri : Their influence on adherence of E . coli to epithelial cells and inflammatory response. *Carbohydrate Polymers*, *141*, 10–19. https://doi.org/10.1016/j.carbpol.2015.12.037

Liu, X. T., Hou, C. L., Zhang, J., Zeng, X. F., & Qiao, S. Y. (2014). Fermentation conditions influence the fatty acid composition of the membranes of Lactobacillus reuteri I5007 and its survival following freeze-drying. *Letters in Applied Microbiology*, *59*, 398–403. https://doi.org/10.1111/lam.12292

Oliveira, A. P., Nielsen, J., & Förster, J. (2005). Modeling Lactococcus lactis using a genome-scale flux model. *BMC Microbiology*, *5*(1). https://doi.org/10.1186/1471-2180-5-39

Teusink, B., Wiersma, A., Molenaar, D., Francke, C., De Vos, W. M., Siezen, R. J., & Smid, E. J. (2006). Analysis of growth of Lactobacillus plantarum WCFS1 on a complex medium using a genome-scale metabolic model. *Journal of Biological Chemistry*, *281*(52), 40041–40048. https://doi.org/10.1074/jbc.M606263200

Walter, J., Loach, D. M., Alqumber, M., Rockel, C., Hermann, C., Pfitzenmaier, M., & Tannock, G. W. (2007). D-Alanyl ester depletion of teichoic acids in Lactobacillus reuteri 100-23 results in impaired colonization of the mouse gastrointestinal tract. *Environmental Microbiology*, *9*(7), 1750–1760. https://doi.org/10.1111/j.1462-2920.2007.01292.x
